# The activity, divergence, and evolutionary degradation of modern-day homing endonucleases and their reconstructed ancestors

**DOI:** 10.64898/2026.07.13.738334

**Authors:** Juliana C. Young, Abigail R. Lambert, Janet M. Young, Lindsey A. Doyle, Miriam Silverstein, David R. Edgell, Barry L. Stoddard

## Abstract

Homing endonucleases (HEs) are selfish genetic elements that drive the mobilization of their own coding sequences, often in concert with surrounding introns. Homing endonuclease genes (HEGs) usually display life cycles in which they accumulate inactivating mutations after invading a host genomic target site, leading to eventual removal from the genome. We identified several hundred novel HEGs and determined the distribution of their proteins’ behaviors and activities. Approximately 10% are expressed as properly folded functional proteins that cleave predictable DNA target sites. Another ∼20% display significant expression but little to no cleavage activity; the remainder display severely reduced expression. Despite the presence of debilitating mutations throughout most HEGs, ancestral reconstructions yielded endonucleases with improved expression and stability. One such reconstruction, at a hypothetical node preceding highly diverged HEs that cleave unique target sites, binds (but does not cleave) their individual targets. It instead cleaves a DNA sequence that represents a hybrid of those modern-day DNA targets, while displaying a specificity profile that resembled those of previously characterized HEs. Its DNA-bound crystal structure adds detail to our understanding of how homing endonuclease DNA contacting surfaces and residues shift and rearrange during evolution, ultimately leading to their action at new target sites.

## INTRODUCTION

Mobile genetic elements (MGEs) are pervasive agents of genomic change and innovation that are found throughout all domains of life (1). Such sequences are typified by DNA transposons (2), retrotransposons (3), plasmids (4), self-splicing introns and inteins (5), integrated and conjugative elements (ICEs) (6), bacteriophage-like elements (7), integrons (8), CRISPR-associated transposons (9), and many other specialized mobile elements. Collectively, they facilitate horizontal gene transfer, genomic rearrangement, and the spread of genetic novelty between species across both single-cell microbial (10–12) and multi-cellular (13) organisms. Regardless of their origin and genomic environment, MGEs often provide little or no selective advantage to their hosts and usually succumb to evolutionary degradation and inactivation due to the effect of accumulated mutations and the action of genomic defense mechanisms (14,15). This dynamic process reflects the balance between a potential benefit to the host (such as the delivery by the MGE of a useful new gene product such as a viral restriction or antibiotic resistance factor) versus a potential cost (for example, insertional mutagenesis leading to inactivation of a host gene, and/or an increase in genome size leading to increases DNA replication time (16)).

Homing endonucleases (HEs), also termed ‘meganucleases’, are highly specific DNA-cleaving enzymes that are encoded either by stand-alone genes or by reading frames embedded within surrounding self-splicing introns or inteins (reviewed in (17)). They are ubiquitous throughout all forms of microbial life, being encoded in phage, bacteria, archaea, and the organellar genomes of single-cell eukaryotes.

Homing endonuclease genes (HEGs) encode proteins that introduce DNA double strand breaks or nicks at specific genomic targets, which are then repaired through homology-driven recombination and the corresponding insertion of the entire mobile element into the site of DNA repair (18–20). When the HEG is a free-standing reading frame, its mobility is generally limited to invasion of intergenic target sites (to avoid disruption of a surrounding host gene (21)). In contrast, when a HEG is embedded within a self-splicing intron or intein, its range of insertion sites is greatly increased, because the entire element can invade host genes without disrupting their expression and the generation of their corresponding products.

The propagation of homing endonuclease genes is extremely efficient, leading to dominant, non-Mendelian inheritance in diploid genomes (18), gene transfer between phage and cellular genomes (22,23), genetic competition between HEGs during mixed phage infections (24), and intron transfer between unrelated subcellular compartments (usually chloroplast and mitochondrial genomes) (25). The accumulation of homing endonucleases and mobile introns in individual genomes can be extremely high. For example, over 10% of the T4 phage genome encodes a total of 12 free-standing and 3 intron-encoded homing endonuclease genes (reviewed in (26)). Similarly, alignment and comparisons of microbial ribosomal DNA sequences indicate the presence of many intervening HEG sequences (27,28).

At least six families of homing endonucleases have been identified. The most well-studied (and the focus of this study) are LAGLIDADG homing endonucleases (‘LHEs’), which contain a small, highly conserved protein domain (typically ∼150 residues in size) encompassing both an extended DNA-binding surface and the enzyme’s active site. They are encoded within fungal mitochondria, algal chloroplast, and archaeal genomes. LHEs exist both as homodimeric proteins (in which two identical protein subunits, each containing a single identical copy of the endonuclease fold, collaborate to act at highly symmetric DNA target sites) and monomeric proteins (in which two non-identical domains are present within a single larger protein chain, allowing them to act at far more asymmetric DNA target sites (17)). LHEs are the most sequence-specific class of HEs, typically recognizing 22 base pair target sites, and are the focus of engineering efforts for a variety of gene editing applications (29–31).

Comparative analysis of a small clade of well-characterized mobile introns and their corresponding LAGLIDADG homing endonuclease genes (all encoding closely related homologues of the I-SceI homing endonuclease in yeast) indicated that such elements display an evolutionary life cycle of invasion, mutational degradation, loss from the genome, and subsequent re-invasion by a related element, with an estimated periodicity of approximately 2 million years (32,33). A focused examination of a single LAGLIDADG endonuclease (I-AniI, encoded in the fungus *Aspergillus nidulans*) and its corresponding DNA target site provided further details of the first steps of the evolutionary degradation of its activity: the enzyme harbors at least two point mutations that have significantly reduced its activity (34), while its insertion target site in the host genome differs from the enzyme’s optimal DNA recognition sequence by at least one basepair (35). The loss of form and function for any particular LHE is not inevitable, however: the acquisition of a secondary “moonlighting” function of importance to the host (for example, their occasional participation in folding and splicing of their surrounding cognate intron, termed ‘maturase activity) can result in host selective pressure to maintain protein folding and stability, which in turn can stabilize their original endonuclease function (36).

For the majority of LHEs whose sole function is to act only as an endonuclease that drives genetic mobility, one would predict that at any moment in time, a significant fraction of current-day homing endonuclease genes would encode dysfunctional proteins, destined for eventual removal from the genome. One study that attempted to test that prediction examined the folding and function of a small group of 34 putative LAGLIDADG homing endonucleases (that all display significant homology to the well-characterized and highly active I-OnuI enzyme). That analysis found that approximately one-third displayed poor folding behavior, versus approximately two-thirds that were capable of cleaving target sites in their cognate host genomes (29).

To assess the same prediction more thoroughly and in a less biased manner (a goal that was motivated and facilitated by the rapidly growing number of LHE sequences that could be identified within microbial and metagenomic sequence databases) we identified a diverse subset of over 300 reading frames corresponding to putative monomeric LAGLIDADG homing endonucleases, all of which are embedded within surrounding group I introns, and then characterized their folding, expression and DNA cleavage activity. Our analysis was focused on monomeric LHEs for three reasons: their predicted structural similarity to several monomeric LHEs that have been previously characterized (17,37), their ability to recognize and act at completely asymmetric DNA target sites, and our own ability to predict their target sites (which correspond to a fusion of the DNA sequences that immediately flank the insertion site of the mobile intron or intein and its corresponding HEG).

Our assessment of the expression and DNA cleavage activity of those candidate enzymes revealed a broad spectrum of behaviors, ranging from well-expressed and fully active endonucleases to those that display significant disruptions in their form and function. Subsequently, we identified additional homologues to generate an even larger collection of over 600 putative LHE sequences, which enabled the creation of a more densely populated phylogenetic tree and the generation of reconstructed ancestral sequences at nodes that precede several different modern-day “extant” homing endonucleases. This allowed us to address two additional questions: (1) whether ancestral reconstruction attempts can generate active endonucleases, despite the fact that their current day descendants are not under strong selective pressure to maintain function and often contain debilitating mutations; and (2) whether the behavior of ancestral enzymes might provide additional insight into the evolutionary divergence of their DNA target site specificity.

## Methods

### Bioinformatic LHE sequence screening

An initial set of 371 sequences encoding putative monomeric LHEs were identified in a 2015 bioinformatic search of all fully sequenced plastid and mitochondrial genomes that were available at that time in the GenBank database (38) for all reading frames that were annotated as containing a group I intron. The exon-intron boundaries for each reading frame were identified, and the collected sequences were examined using a Psi-BLAST search (39,40) to identify potential LHEs. All candidates were further culled based on a cutoff of 20% or higher sequence identity to any one of nine query sequences available at that time with known, validated endonuclease activity against established target sequences and corresponding DNA-bound structures (DNA-bound crystal structures: I-AabMI PDB ID 4YIT, I-CpaMI 4YIS, I-GpeMI 4YHX, I-GzeII 4Z1X, I-LtrI 3R7P, I-LtrWI 4LQ0, I-OnuI 3QQY,I-PanMI 5ESP, and I-SmaMI 5E5O). Putative ORFs shorter than 200 amino acids were assumed to exist as homodimers and were excluded.

Several years after our initial search (in 2023), we expanded our initial collection of LHE sequences via an additional search of available sequence databases, and identified an additional 246 putative LHE genes. This larger set of 625 coding sequences was used for to select subsets of sequences to use for computational ancestral sequence reconstruction efforts.

### Cloning of synthetic LHE ORFs

A small “pilot” set of 18 putative LHE ORFs from our original bioinformatic search were first ordered from Genscript (Piscataway, NJ) as synthetic genes cloned into the pETCON vector, for expression on the surface of yeast. Those constructs are denoted hereafter with identifiers corresponding to “HE##” throughout this document, corresponding figures, and ***Supplementary Information***. The remaining 353 LHE sequences (which are denoted hereafter with identifiers corresponding to “Gen9-###”) from the 2015 dataset were ordered from Gen9 (Cambridge, MA) as synthetic genes delivered in their pG9m-2 plasmid. Those constructs were synthesized commercially via a parallel high-throughput approach at Gen9 Bio (relying on a technology previously described in (41)) and then sequence-verified. 30 sequences were not able to be synthesized from the Gen9 order, leaving a total of 341 LHE ORFs that were delivered.

The synthetic LHE ORFs from Gen9 were subcloned *en masse* into the pETCON yeast surface display vector (Addgene #41522; (42)) via homologous recombination in yeast, and later isolated as single clones. First, 100 ng of each synthetic plasmid was mixed to create a pool of PCR templates. This pool of LHE ORFs was PCR-amplified using primers designed to create 80-90 base pair overlaps with each end of a linearized pETCON yeast surface display vector (generated via digests with NdeI and XhoI restriction enzymes). The pool of PCR-amplified LHE inserts was transformed into EBY100 yeast (American Type Culture Collection, ATCC) alongside open pETCON vector using a lithium acetate transformation protocol (43). The yeast transformation reaction was cultured overnight, and plasmids containing cloned LHE sequences were extracted from the yeast using a 2-hour Zymolase (Zymo Research) digest followed by a modified plasmid extraction protocol utilizing Qiagen miniprep kit components (Qiagen). While cloning in batch format was most effective using yeast, the necessary plasmid prep and sequence verification steps were more efficiently performed in bacteria. Therefore, the plasmids from the yeast miniprep were transformed into DH5α *E. coli* (New England Biolabs) and plated. Individual bacterial colonies were sequenced to identify successfully cloned LHE ORFs. Plasmids were isolated from bacterial cultures containing unique LHEs, and these plasmids were transformed back into yeast for further analysis. This process was repeated until clones were isolated for 307 unique LHEs (34 LHE sequences were set aside after multiple attempts at subcloning failed).

### Yeast induction of surface expression

Yeast cultures were grown from single colonies at 30°C with shaking in selective culture media (Synthetic complete or ‘SC’ media, -Ura -Trp, Sunrise Science Products) + 4% glucose (the higher percentage of glucose over the typical 2% helps to discourage flocculation). For induction of surface expression, yeast cultures were grown from a starting density of 24 million/mL in SC + 2% raffinose + 0.1% glucose for 6 hours at 30°C with shaking, washed with sterile water, and then incubated at a density of 20 million/mL in SC + 2% galactose overnight on the benchtop (room temperature).

### Yeast staining for surface expression

LHEs expressed on the surface of yeast were analyzed by flow cytometry as previously described (37). Briefly, yeast were stained for 45-60 minutes at 4°C in a yeast staining buffer (10 mM HEPES pH 7.5, 180 mM KCl, 10 mM NaCl, 0.1% w/v galactose, 0.1% w/v bovine serum albumin) with an anti-Myc-FITC antibody (ICL) along with either an anti-HA-APC antibody (BioLegend) or an anti-HA-biotin/Streptavidin-phycoerythrin (SAV-PE) conjugation (BioLegend and Thermo Fisher Scientific) (**Figure 1a**). Full-length expression of the LHE was indicated by the presence of FITC signal after staining of the C-terminal Myc tag. Staining of the N-terminal HA tag with a second color could be used to identify LHEs that were not expressed all the way through to the C-terminal Myc tag (presumed to be stalled, unstable, or otherwise truncated).

**Figure 1.**
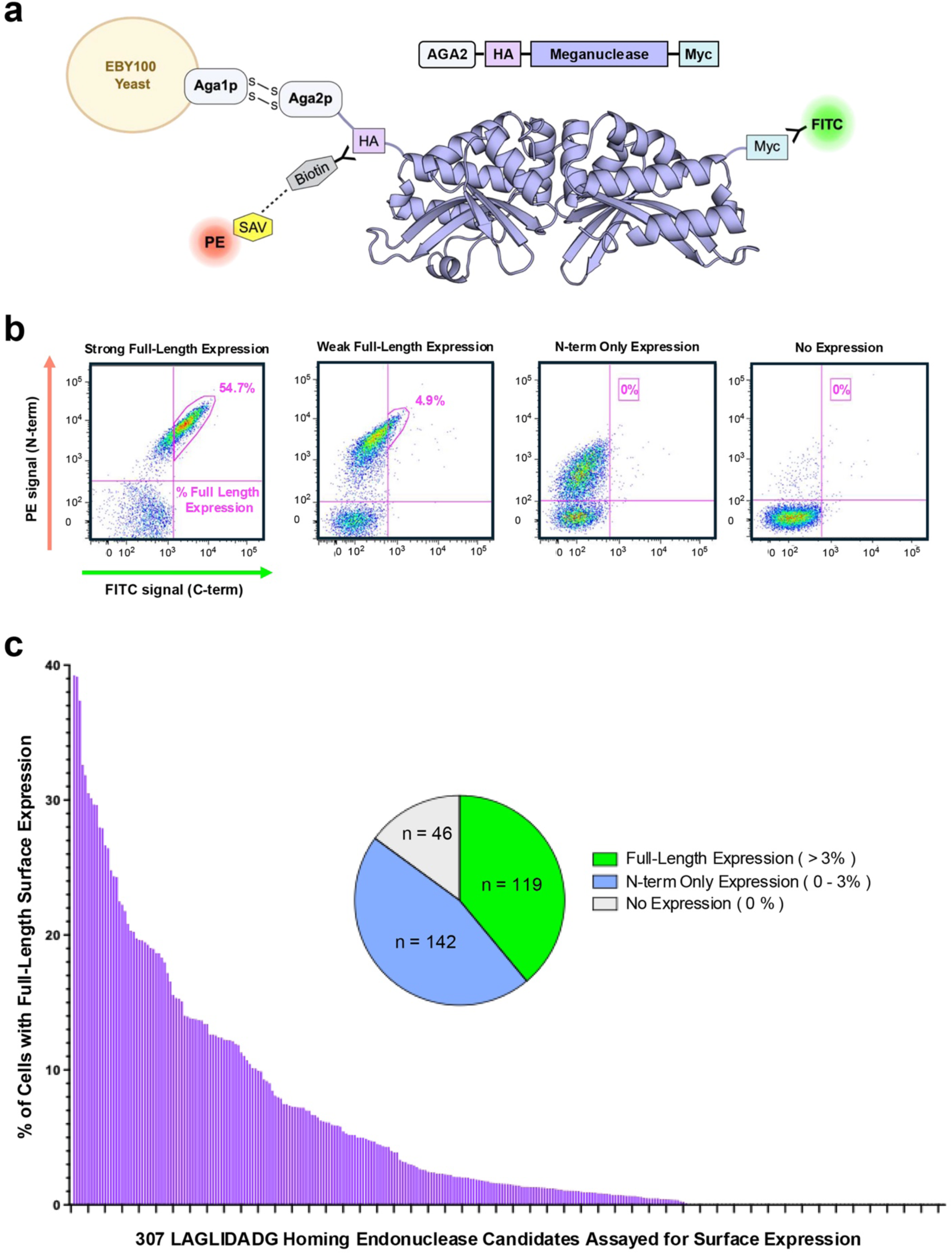
Distribution of LHE expression levels. **Panel a:** Schematic of LAGLIDADG homing endonuclease (LHE) expression on the surface of yeast. The pETCON yeast surface display vector expresses the cloned LHE with an N-terminal hemagglutinin (HA) tag, a C-terminal Myc tag, and as a fusion to the secreted Aga2p yeast cell surface binding protein. The disulfide-mediated interaction between the Aga2p fusion protein and the Aga1p protein partner on the exterior of the EBY100 yeast cell anchors the expressed LHE to the yeast surface. We stain the N-terminal tag using an anti-HA-biotin antibody conjugated to a fluorescent streptavidin protein (SAV-PE), and the C-terminal tag using an anti-Myc fluorescent (FITC) antibody. Signal from both tags indicates full length protein successfully expressed on the surface of yeast. **Panel b:** Quantification of surface expression was performed in FlowJo by drawing gates around the cells in each sample with positive FITC signal (after staining of the C-terminal Myc tag with an anti-Myc-FITC antibody) and quantifying the percentage of the total cells which fall within that gate. Four sample flow cytometric plots show examples of quantification for strong full-length expression, weak full-length expression, N-term only expression, and no expression. For quantification of N-term only and no expression samples, a small rectangular gate was drawn in an area of the graph with no noise to ensure that the quantification gate would read a value of zero. **Panel c:** Quantified surface expression for the collection of 307 candidate LHEs. Each bar represents the percent of full-length surface expression for one characterized LHE, ordered along the x-axis from highest surface expression to lowest. **Inset**: Pie chart representation of LHEs with full-length expression (expression of greater than 3%, n = 119), N-term only expression (expression of 0 – 3%, n = 142) and no expression (n = 46). See the **Deposited Raw Data** for the flow plots, quantification gates, and corresponding values used to create this graph.

These experiments were performed in batches over many months, so we chose to quantify and compare levels of surface expression between LHEs using the overall percentage of cells with FITC signal rather than comparing the magnitude of the FITC signal (which varies depending on factors like expression level on a given day and voltage settings on the instrument). Flow cytometric data was analyzed with FlowJo (44) and the percentage of cells with full-length surface expression was quantified by drawing gates around populations of cells with detectable FITC signal (**Figure 1b**). Fluorescent signal from staining of the N-terminal HA tag was not quantified. An average percent expression value was calculated for those constructs which were assayed multiple times (see **Deposited Raw Data**).

### Generation of labeled DNA target site substrates

Double-stranded DNA oligonucleotide target site substrates were generated as previously described (45). Briefly, an oligo template containing a 58 bp DNA target region flanked by two 16 bp primer binding sequences was amplified by PCR with Platinum Taq High Fidelity DNA polymerase (Invitrogen) using labeled primers which add a biotin to one end of the PCR product and an AlexaFluor647 (A647) fluorophore to the opposite end (IDT DNA). The A647 label can be added to either the 5’ or the 3’ end of the DNA target substrate. The resulting dual-labeled, double-stranded PCR products were digested for 6 hours with Exonuclease I (New England Biolabs) to remove any unincorporated primers or leftover single-stranded template. The double-stranded DNA target substrates were then purified using a small G-100 Sephadex size exclusion column (GE Healthcare) in a filter plate.

The position of the A647 label (on either the 5’ end or the 3’ end of the DNA target substrate) does not affect the output of non-tethered cleavage assays. However, we typically make and test substrates with both possible substrate labeling schemes (A647 on the 5’ end vs. A647 on the 3’ end) for the tethered flow cytometric cleavage assay, since some LHEs remain bound to one end of the cleaved DNA more tightly than the other. This end-holding behavior can affect the magnitude of the drop in A647 signal which indicates successful cleavage activity in the tethered cleavage assay. The A647 label was added to the 5’ end of the DNA target substrates used in this study.

### DNA cleavage assays using surface-displayed endonucleases (“tethered” DNA cleavage assays)

LHEs expressed on the surface of yeast were analyzed for DNA cleavage activity using a previously described tethered flow cytometric cleavage assay (37). Briefly, the C-terminal Myc tag was stained with anti-Myc-FITC antibody, and double-stranded dual-labeled DNA target site substrate was physically tethered to the N-terminus of each surface-expressed LHE (2 million yeast cells per well of the experiment) using an anti-HA-biotin/Streptavidin-PE (SAV-PE) conjugation (40 nM DNA target substrate and 5 nM SAV-PE) (**Figure 2a**). The stained yeast were divided into two duplicate wells and incubated in 50 microliters cleavage buffer (10 mM HEPES pH 7.4, 150 mM KCl, 10 mM NaCl, 5 mM K-glutamate, 0.05% bovine serum albumin) for 30-60 minutes at 37°C in the presence of 5 mM MgCl_2_ (allows for cleavage to occur) or 5 mM CaCl_2_ (cleavage of the tethered DNA substrate can NOT occur). The yeast were analyzed by flow cytometry to measure the A647 signal from the cleaved vs. uncleaved tethered DNA target substrate in the magnesium vs. calcium samples. Successful target cleavage leads to a drop in A647 signal when the fluorescent tag on the DNA substrate is separated and washed away (**Supplementary Figure S1**).

**Figure 2.**
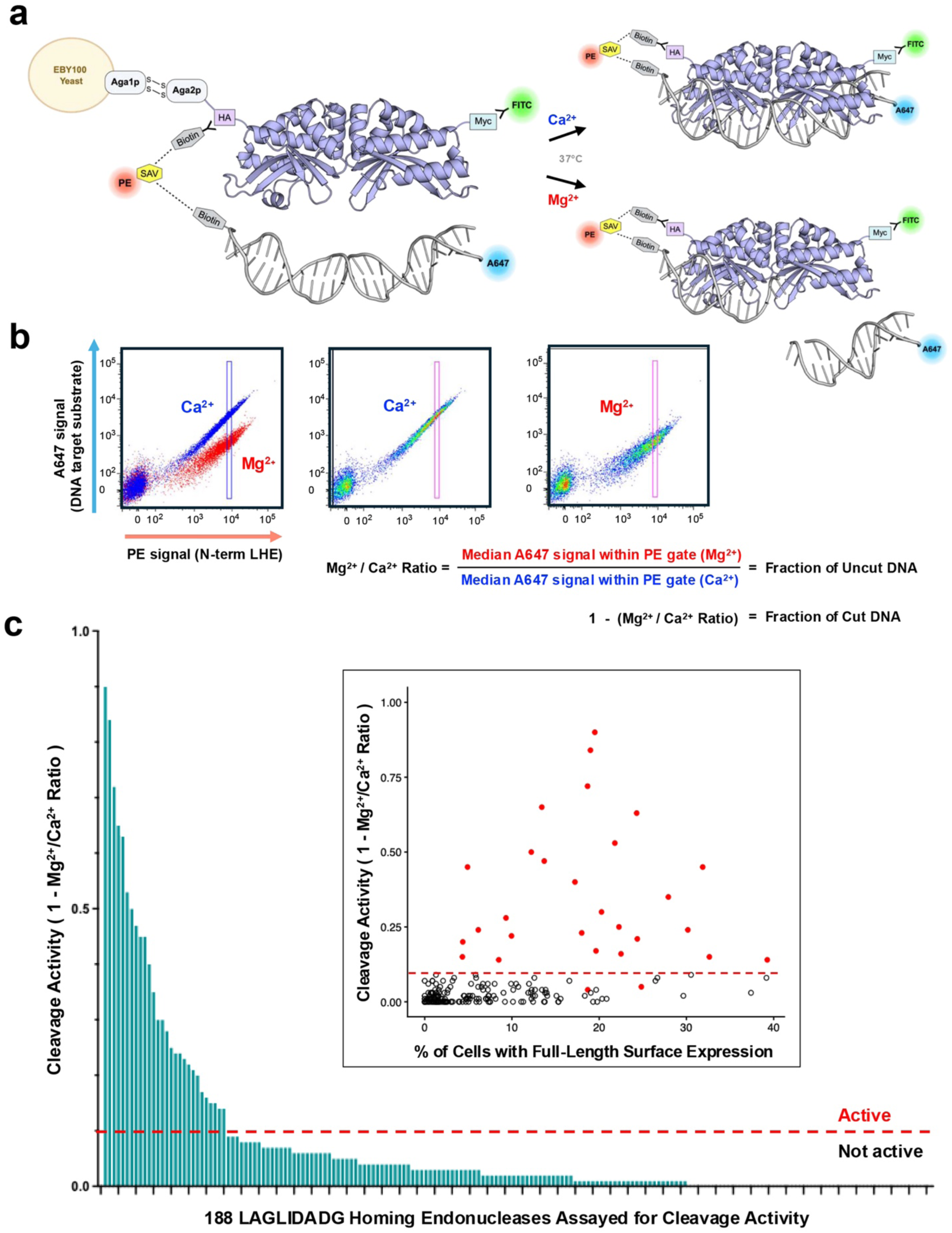
Distribution of LHE Cleavage activities. **Panel a:** Schematic of the tethered flow cytometric DNA cleavage assay. The C-terminal Myc tag is stained with anti-Myc-FITC antibody for visualization of full-length protein expressed on the surface of yeast, and a double-stranded DNA target substrate labeled with both biotin and Alexa-647 is tethered to the N-terminal HA tag of the surface-expressed LHE through a biotin-streptavidin bridge. The cells are divided into two duplicate wells and incubated at 37°C in a digest buffer with either calcium (Ca^2+^) or magnesium (Mg^2+^). In the presence of calcium, the LHE cannot cleave its target; in the presence of magnesium, the LHE can cleave its target, resulting in separation and dissociation of the A647 tag from the tethered complex in a subsequent wash step. **Panel b:** Quantification of DNA cleavage activity in the tethered flow cytometric assay. The tethered DNA target substrate in the presence of calcium (blue population of cells) produces a diagonally shaped population when plotting the signal from the A647 tag on the DNA target substrate (y-axis) vs. the signal from the streptavidin-phycoerythrin (SAV-PE)-biotin staining of the N-terminal HA tag (x-axis). Cleavage of the DNA target in the presence of magnesium results in a drop in A647 signal (red population of cells) when the A647 tag is washed away. DNA cleavage activity in this assay was quantified by calculating the median A647 signal from the tethered DNA target substrate in the presence of calcium vs. magnesium within a thin rectangular gate along the PE axis. This median A647 value presented as a ratio of Mg^2+^/Ca^2+^ represents the fraction of uncut DNA target substrate remaining in the sample. Therefore, a value of 1 – (Mg^2+^/Ca^2+^) is equal to the fraction of cleaved substrate. **Panel c:** Quantified cleavage activity for 188 characterized LHEs. Each bar represents the cleavage activity (fraction of cleaved DNA target substrate, or 1 - (Mg^2+^/Ca^2+^ Ratio)) for one characterized LHE, ordered along the x-axis from highest cleavage activity to lowest. A dotted red line marks a cleavage value of 0.1, which serves as the threshold for an LHE to be considered “active” in this assay. (**Inset**) Cleavage activity visualized as a plot of 1 - (Mg^2+^/Ca^2+^ Ratio) (y-axis) vs. % of cells with full-length surface expression (x-axis), with 29 total active enzymes designated as red dots and inactive enzymes designated as open black circles. Two red dots fall below the threshold value of 0.1, as they were observed to be active in the complementary non-tethered in vitro cleavage assay. See the **Deposited Raw Data** for the flow plots, quantification gates, and corresponding values used to create these graphs.

This DNA cleavage activity can be quantified by drawing a thin rectangular gate along the PE axis where the cell population is most dense and calculating the “Mg^2+^/Ca^2+^ ratio” within that gate, equal to the ratio of the median A647 signal from the magnesium (cleavage) sample to the median A647 signal from the calcium (no cleavage) sample (**Figure 2b**). The presence of A647 signal in this experiment is due to uncleaved tethered DNA target substrate, so the Mg^2+^/Ca^2+^ ratio represents the fraction of uncleaved DNA target substrate remaining. Therefore, the value of 1 – (Mg^2+^/Ca^2+^ ratio) indicates the fraction of tethered DNA target substrate that was successfully cleaved, and this is the value chosen to represent DNA cleavage activity plotted in **Figure 2c**. A sample with no cleavage of the tethered substrate (i.e. no change in A647 signal) has a Mg^2+^/Ca^2+^ ratio of 1, and the corresponding value of 1 – (Mg^2+^/Ca^2+^ ratio) equals zero. See the **Deposited Raw Data** and **Supplementary Information** for flow cytometric plots and a table of quantified cleavage values, respectively.

### DNA cleavage assays using endonucleases released from the yeast surface (“untethered” DNA cleavage assays)

Approximately 5 million yeast cells with surface-expressed LHE were incubated for 30-60 minutes at 37°C in 40 microliters cleavage buffer (described above) along with 5 mM MgCl_2_, 20 nM A647-labeled DNA target site substrate, and 10 mM dithiothreitol (DTT). The DTT reduces the disulfide bonds of the AGA1-AGA2 interaction that anchors the expressed LHE to the yeast surface, leading to the release of the enzyme into solution. Depending on the strength of expression, the resulting free protein concentration is estimated to range from roughly 0.3 to 3 nM per 1 million expressing yeast cells in 50 microliters of volume (46). A colorless Ficoll loading dye was added to the reactions before loading samples onto an 11.3% acrylamide gel. The gel was run at 120 volts for 60 minutes, and cleavage bands were visualized using the Cy5 channel on a Typhoon fluorescent scanner (Amersham/GE) (**Supplementary Figure S2a**).

### Determination of site and centering of DNA target cleavage via run-off DNA sequencing

For the characterization of the 29 active enzymes identified in the first part of this study, 5-10 million yeast cells with surface-expressed LHE were incubated for 1-2 hours at 37°C with approximately 1 microgram (∼ 2 nM) of linearized target site plasmid in 50 microliters of cleavage buffer containing 10 mM dithiothreitol (DTT) and 5 mM MgCl_2_. For later experiments during the characterization of ancestral reconstructions, the digest was performed with 100 nM purified recombinant protein combined with circular target site plasmid at a similar (∼ 2 nM) substrate concentration. In either case, the entire cleavage reaction was loaded onto a 1% agarose gel to separate the cleaved bands. Cleavage bands (either two cleaved fragments from a linear DNA target or a single linearized band after cleaving circular plasmid DNA) were extracted from the gel and purified with the Zymoclean Gel DNA Recovery Kit (Zymo Research). Purified bands were used as templates for individual BigDye (Thermo Fisher Scientific) Sanger sequencing reactions.

The resulting run-off sequencing chromatograms revealed a single large adenine peak where the Taq polymerase came to the end of the cleaved template. This large “A” peak marked the precise position where that strand of DNA was cleaved by the LHE. In cases where two cleavage product bands were not able to be purified, a single clear run-off sequencing reaction was sufficient to determine the center of the DNA target site given the verified characteristic that LAGLIDADG LHEs cleave their targets to create 4-base 3’ overhangs. In the absence of sufficient purified cleavage products or clean run-off sequencing data, the center of some target sites could still be determined through alignments with known target sites from closely related LHEs. See the **Deposited Raw Data** for the chromatograms produced from run-off sequencing reactions and their corresponding analysis.

### Ancestral sequence reconstructions

In order to reconstruct selected ancestral sequences in the I-OnuI (focal sequences I-OnuI and I-GpeMI) and I-PnoMI groups (focal sequences PnoMI, HE08 and Gen9-348), we first used an expanded collection of 625 homing endonuclease sequences to identify members of each group as follows.

We constructed a tree of all LAGLIDADG domains in our collection of 625 sequences (**Supplementary Figure S4**). Each monomeric LHE usually contains two such domains: we used the hmmsearch tool (HMMER package version 3.3.2) with the “LAGLIDADG_1” Hidden Markov Model (PFAM PF00961.22) (47) as query to extract all domain sequences in the form of an alignment to the HMM. We filtered this alignment to remove sequences with an ungapped length of <75aa (the HMM has 102aa positions), and to remove alignment positions where >=75% of sequences contained a gap. This filtered alignment contained 1140 domain sequences and 102aa positions. We used ProtTest (version 3.4.2) (48) to infer that best fitting amino acid substitution model is the CpREV model. We then used PHYML (version 3.3.20200621) (49) to estimate a phylogeny, estimating the proportion of invariable sites, with 4 rate categories and estimating the shape of the gamma distribution (i.e. parameters “--pinv e --alpha e -f e -m CpREV”).

We analyzed the resulting LAGLIDADG domain phylogeny using R (50) with the ape (51) and ggtree (52) packages. For each sequence group of interest, for each of the two LAGLIDADG domains, we identified the MRCA (most recent common ancestor) of the group’s focal sequences, then stepped back three nodes deeper in the tree in order to obtain an extended clade that includes outgroups. We selected all descendent tips of these “MRCA-minus-3” nodes, merged the lists obtained from the two LAGLIDADG domains of all focal sequences in each group, and extracted any HEs with full-length sequences (>=280aa) for use in ancestral reconstruction.

For each group, we then aligned full-length selected sequences using MAFFT (version 7.453) (53), trimmed the alignment to remove regions outside the extent of the focal sequences, and removed any sequences whose ungapped length was now <250aa. We also generated a non-redundant alignment, because some sequences were identical to one another within the trimmed region. We also noticed that our sequence list for the I-OnuI group included three engineered sequences, which we removed from further analysis (ancestral reconstruction methods assume sequences evolved naturally). The resulting alignments contained 40 (I-PnoMI clade) sequences or 18 (I-Onu-I clade) sequences (**Figure 3**).

**Figure 3.**
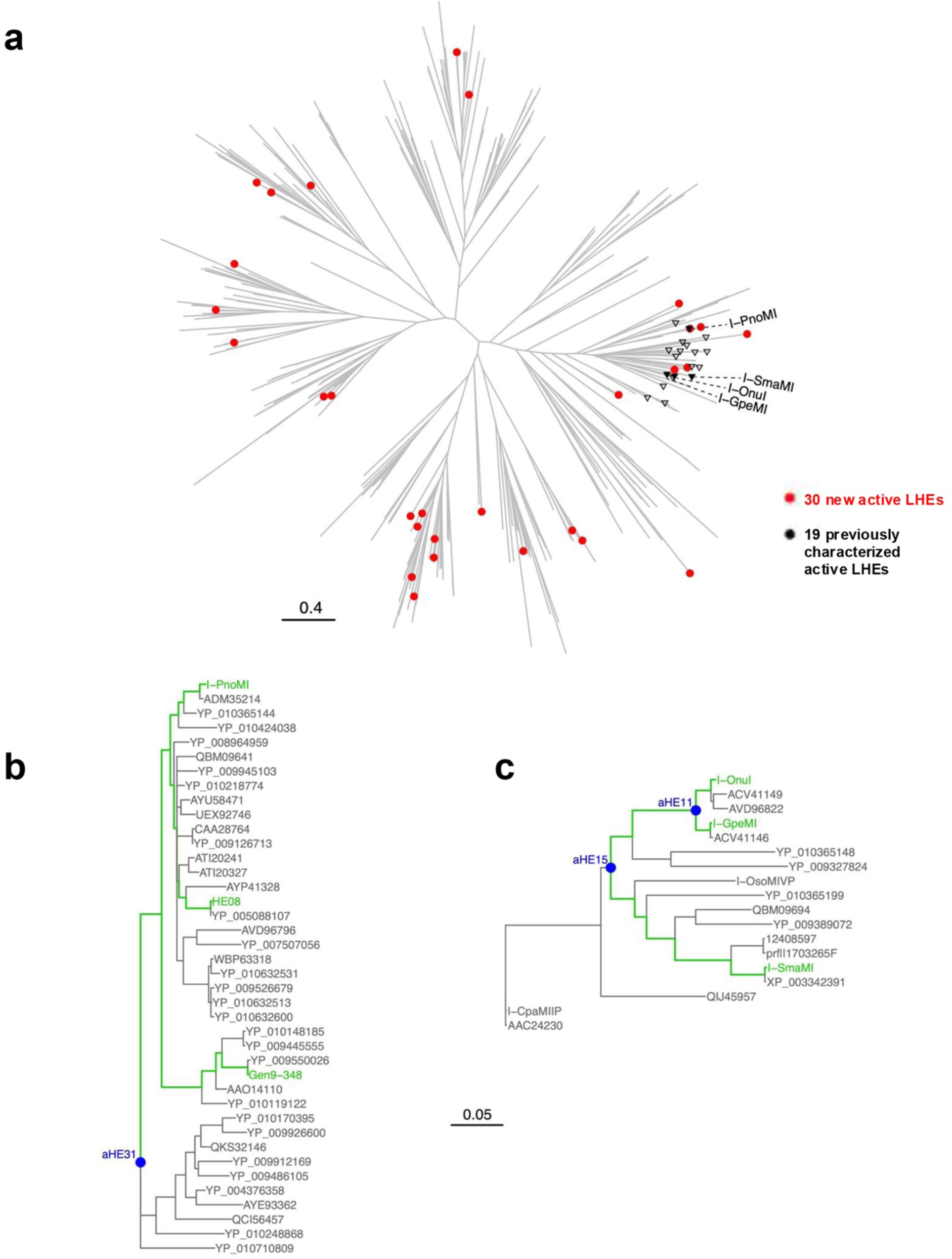
LHE phylogenetic trees. **Panel a**: Phylogenetic tree generated from the set of 19 previously characterized enzymes (black triangles) plus 371 candidate LHEs identified by the 2015 bioinformatics search. Red circles designate the 29 active LHEs identified by this study, which span a much greater range of diversity than the 19 previously characterized enzymes. The scale bar indicates number of amino acid substitutions per site. Filled black triangles indicate the labeled enzymes, which comprise the focal sequences for our ancestral reconstruction studies. **Panel b:** Ancestral reconstruction in the I-PnoMI clade. Using a domain-based tree created from our larger set of 622 LHEs (Supplementary Figure S4), we selected LHE sequences from a subclade containing three closely related, validated LHEs with known target sites (I-PnoMI, Gen9-348, and HE08: green labels), as well as selected outgroups (see Methods). We used the BaliPHY algorithm to estimate a consensus tree of this subset of sequences and to reconstruct ancestor aHE31 (labeled in blue). **Panel c:** Ancestral reconstruction in the I-OnuI clade. As for the I-PnoMI clade, we selected sequences from the subclade containing two closely related LHEs (I-OnuI and I-GpeMI), as well as selected outgroups, including another previously characterized enzyme, I-SmaI. We first characterized aHE11, a recent ancestor of I-OnuI and I-GpeMI, before extending our study to a deeper (older) node, aHE15. Panels b and c are plotted on the same scale; the scale bar applies to both panels.

Next, we performed ancestral reconstruction on each trimmed alignment, using the BaliPhy suite (version 3.6.0) (54). In more detail, we used the bali-phy command to run two separate Markov chains to simultaneously estimate phylogenies and alignments. We ran each chain overnight, by which time all chains had gone through >27,000 iterations, and used the bp-analyze command to summarize chain statistics. We used trees-consensus and tree-tool to determine the consensus tree over both chains and ran the tree-tool command again to add internal node names to the consensus tree. We used the cut-range, alignment-chop-internal and alignment-max commands to extract the most likely alignment of extant sequences, skipping the first 1000 samples in each chain. We then used the summarize-ancestors command to add inferred ancestral sequences to the summary alignment.

For each clade, we inspected the resulting consensus trees and summary alignments, and selected ancestral nodes to test experimentally. For ancestors in the PnoMI clade, we tested Bali-Phy’s ancestral reconstructions without modification. For ancestors in the I-OnuI clade, we noticed that indel positions often contained amino acid predictions in the ancestral reconstructions, even when a gap seemed a more likely ancestral state according to parsimony principles. We created edited versions of the ancestral reconstructions that contain gaps at these positions, and proceeded with experimental testing.

### Expression and purification of recombinant LHEs

LHEs were expressed as untagged constructs in the pET21d bacterial expression vector (Novagen) using either *Escherichia coli* BL21-CodonPlus (DE3)-RIL (Agilent) or Rosetta (DE3) (Novagen) competent cells. Both cell types were cultured in 1 L Luria Broth (LB) media (RPI) with 100 ug/mL of ampicillin. Each liter of LB-amp was inoculated with 10 mL of a stationary phase overnight culture and grown at 37°C with shaking until reaching an OD_600_ of 0.6-0.8.

Expression was induced by adding IPTG to a final concentration of 0.5 mM, and the cultures were placed in a shaking incubator at 16°C overnight. Each liter of culture was then pelleted and stored at -20°C until ready for purification.

Recombinant LHEs were purified in two stages, using heparin ion exchange chromatography followed by size exclusion chromatography. Frozen bacterial expression pellets were thawed and resuspended in buffer (25 mM Tris-HCl pH 7.5, 200 mM NaCl) supplemented with 50 micromolar PMSF and 0.01 U/μL benzonase nuclease. The cells were broken open using sonication, and the resulting cell debris was pelleted by centrifugation for 20 minutes at 18,000 RPM. The supernatant was removed and filtered through a 0.45 micron syringe filter before loading onto a 5 mL Heparin HP HiTrap Column (Sigma-Aldrich). A linear salt gradient was used to elute the bound LHE, and fractions containing the correct sized protein were pooled and concentrated to a volume of 5 mL or less. The concentrated protein sample was filtered through a 0.22 micron pore size filter and loaded onto a Superdex HiLoad 16/60 prep grade size exclusion column (Sigma-Aldrich) using a 5 mL loop size. The SEC buffer used was as follows: 25 mM Tris-HCl pH 7.5, 200 mM NaCl, and 5% glycerol. See **Supplementary Figure S5b** for gels from the purification of aHE31 and **Supplementary Figure S6b** for gels from the purification of aHE15.

### Determination of LHE thermal stability via thermal denaturation and circular dichroism spectroscopy (CD)

Plane polarized light was measured with a J-800 CD spectrometer (JASCO). Purified LHEs were dialyzed into CD buffer (10 mM potassium phosphate, pH 7.0), diluted to 25 micromolar concentration, and scanned in a 1 mm path length cuvette at a wavelength of 210 nm across a range of temperatures from 20°C - 98°C. The melting temperature (T_m_) was calculated for each enzyme using denaturation analysis in Spectra Manager II software provided by JASCO.

### In vitro cleavage assays using purified recombinant LHEs and 500bp target site substrates

*In vitro* cleavage reactions varying either digest time (5 – 60 minutes) or enzyme concentration (12.5 nM – 400 nM) were performed to compare the cleavage activity of purified recombinant ancestor aHE31 to that of related modern-day enzyme I-PnoMI. For all digest reactions, 1.5 µg of purified 500 bp substrate containing the I-PnoMI target site sequence was combined with cleavage buffer at pH 7.4 (described above), 5 mM MgCl_2_, and purified recombinant enzyme in a total reaction volume of 40 μL (corresponding to a DNA substrate concentration of ∼5 nM). Each reaction was incubated at 37°C and stopped with the addition of 4 μL of 0.5 M EDTA solution. Reactions examining the effect of increasing incubation time used an enzyme concentration of 100 nM and varied the total digest time from 5 - 60 minutes. Reactions looking at the effect of increasing enzyme concentration used an incubation time of 30 minutes and varied enzyme concentration from 12.5 nM - 400 nM. Cleavage products were separated on a 1.7% agarose gel at 120V for 30 minutes and visualized on a BioRad Gel Doc imager.

The target site substrate used for this set of reactions was generated through PCR amplification of a 500 bp region of an I-PnoMI target site plasmid using PhusionPlus polymerase (Thermo Fisher Scientific).

The PCR product was purified by gel extraction from a 1% agarose gel using the Zymo Research gel extraction kit and eluted with 30 μL of water. The purified target concentration was determined by measuring the UV absorbance at 260nm with a nanodrop spectrophotometer.

### Yeast SELEX library generation and screening

In order to determine whether ancestral reconstruction aHE15 was an active endonuclease, and if so what its DNA target specificity corresponded to, we carried out a selection experiment in which (1) a random, unbiased DNA target library was panned against surface-displayed aHE15, to select those sequences that could be bound by the enzyme under moderate stringency (300 mM KCl, pH 7.4), followed by (2) all captured DNA sequences that could be bound by the enzyme were screened for their ability to be cleaved by recombinant aHE15. For both steps, the selection process was carried out for multiple rounds to enrich the selected DNA targets for binding and then cleavage. Details are provided below.

The generation of the SELEX library substrate for binding-based selection was performed according to a previously described SELEX protocol (55,56) with a few customized parameters noted here. In brief, aHE15 was induced on the surface of yeast (described above), and 3 million yeast cells were used for each experiment. The double-stranded DNA target substrate for each round of selection was generated by PCR amplification using HiFi Platinum Taq polymerase (Invitrogen) with the initial “SELEX0” fully randomized target created from an oligonucleotide template (IDT) containing 30xN bases flanked by two primer binding sequences (**Supplementary Figure S8a**). Three thymidine bases preceding and three adenine bases following the 30xN sequence were added to discourage high-affinity binding to the constant regions of the substrate. A small amount of A647-labeled SELEX0 substrate was generated for use in a series of optimization tests which varied the concentrations of both KCL and total DNA target substrate in the reaction. For each test condition, 3 million induced yeast cells were incubated for 30 minutes at room temperature in a total volume of 100 μL binding buffer (100 mM NaCl, 10 mM HEPES pH 7.5, 50 mM K-Glu, 2 mM CaCl_2_) plus a range of 200-300 mM KCl and 250-2000 nM labeled SELEX0 DNA substrate. The goal of binding approximately 5% of the initial randomized oligo substrate was achieved with a combination of 300 mM KCl and 1000 nM DNA target substrate (**Supplementary Figure S8b**).

This set of parameters (3 million induced yeast cells, binding buffer with 300 mM KCl, 1000 nM DNA target substrate, 100 μL reaction volume, and 30-minute room temperature incubation) was then applied to multiple rounds of SELEX binding, starting with a fully randomized, non-labelled SELEX0 DNA target substrate. After each incubation step, the yeast were washed six times and resuspended in 10 μL buffer EB (10 mM Tris-HCl, pH 8.5). Next, the yeast were heated in a thermocycler for 10 minutes at 90 °C to release any bound DNA substrate. This elevated incubation temperature was necessary due to the usually high T_m_ of the aHE15 enzyme (78°C), thus requiring a higher temperature than the original published SELEX protocol for complete denaturation of the protein to release the bound DNA substrate. After incubating at 90 degrees, the samples were spun down to pellet the yeast, and the supernatant containing the heat-released DNA was collected. This pool of bound DNA sequences was then used as the template for another set of PCR reactions to generate the double-stranded DNA target substrate for the next round of SELEX (SELEX1). This binding selection process (**Figure 6a**) was repeated for a total of four rounds using the same culture of galactose-induced yeast to ensure equivalent levels of surface-expressed aHE15 for each round.

### In vitro cleavage library generation and screen

The *in vitro* cleavage-based screen was performed largely as described previously (35). The cleavage selection steps were performed using purified recombinant aHE15 protein instead of yeast surface-expressed enzyme, and the DNA target sites were presented in the form of a plasmid library rather than short double-stranded oligonucleotide substrates. The initial target site library was generated from the DNA target substrates collected from Round 4 of the SELEX binding screen. Primers were designed to add 22 bp Gibson overlaps to the DNA target substrates isolated from SELEX4 to allow for cloning into open pIDT-SMART vector (Genscript) using the NEBuilder HiFi DNA Assembly kit (New England Biolabs):

SELEX_Gib_forward: CACTGCTCGATCCGCTCGCACCCAGGGATCCATGCACTGTACGTTT
SELEX_Gib_reverse: GTCGCGCTGCTCTCGTCGATCCAGGATCCGCAGTCAAGTGGTTT

For each cleavage selection step, 50 nM of both the target site library plasmid and recombinant aHE15 enzyme were mixed in a 40 μL volume with cleavage buffer pH 7.4 (described above) and 10 mM MgCl_2_ to facilitate cleavage. The digest was incubated at 37°C for 2 hours, and the entire reaction was loaded into a 0.5% agarose gel and run at 120 volts for 40 minutes. Linearized plasmid was not visible on the gel until Round 3, so both uncut/supercoiled plasmid DNA and a PvuI-linearized plasmid reference were run in adjacent lanes of the gel to help locate the area where the cleaved product should be. This designated area was extracted regardless of whether or not the linearized product was visible. The cleaved/linearized plasmid was extracted and purified using a gel extraction kit (Zymo Research) and eluted with 15 μL of water. The purified linearized plasmid was then re-ligated back into circular DNA using T4 ligase (ThermoFischer Scientific) and transformed into competent DH5-alpha bacterial cells.

Half of the transformation reaction was plated onto LB-AMP plates and half was used to start a 5 mL LB-AMP overnight miniprep culture. 48 colonies from the plates were sequenced by colony PCR (**Supplementary Figure S9**), and a miniprep was performed to generate the plasmid substrate for the next round of cleavage selection. A total of four rounds of cleavage selection were performed (**Figure 6b**).

### In vitro cleavage assays using purified recombinant aHE15 and short A647-labeled target site substrates

Purified recombinant aHE15 was incubated at 100 nM concentration in a 40 μL reaction volume containing cleavage buffer at pH 7.4 (described above) with 20 nM of an A647-labeled double stranded DNA target site substrate (I-OnuI, I-SmaMI, 4xSeq, 17xSeq) and 5 mM divalent cation (CaCl_2_ or MgCl_2_). Digests were performed at 37°C for 60-120 minutes, and the reactions were stopped by adding 4 μL of 0.5M EDTA to 36 μL of the digest and cooling on ice. A colorless Ficoll loading buffer was added to each reaction, and samples were run on an 11.3% acrylamide gel for 60 minutes at 120 volts. Gels were visualized on a Typhoon fluorescent scanner (Amersham/GE).

### One-off specificity profile using short A647-labeled DNA target site substrates and purified recombinant aHE15

Purified recombinant aHE15 protein was assayed against a set of 66 “one-off” DNA target substrates, with each substrate containing a single base change away from the most enriched target sequence found in our cleavage screen (“17xSeq”) in order to identify the preferred DNA base at each position of the DNA target. Target site templates were designed based on the centered 22bp 17xSeq target identified by the cleavage selection screen, synthesized by IDT, and generated as described above. 67 total reactions were run (17xSeq plus the set of 66 one-off substrates), containing 50 nM purified recombinant aHE15 enzyme, 50 nM A647-labeled DNA target substrate, and cleavage buffer at pH 7.4 (described above) supplemented with 5 mM MgCl_2_ in a total volume of 40 μL. The reactions were incubated for 2 hours at 37°C degrees and stopped by adding 10 μL of colorless Ficoll loading dye and moving to ice. 11 μL of each sample was run on an 11.3% acrylamide gel for 60 minutes at 120 volts. Gels were then visualized on a Typhoon fluorescent scanner (Amersham/GE), and cleavage bands were quantified by densitometry using ImageJ software. Cleavage efficiency was quantified by determining the ratio of “cut” / “uncut” using the density of bands for cut and uncut substrate. This value was normalized relative to the ratio of cut / uncut product for the aHE15 enzyme against the original enriched target sequence (17xSeq). A total of six acrylamide gels were required to visualize samples from all one-off target substrates. A duplicate sample of the 17xSeq cleavage reaction was run on each gel, and the densitometry analysis was performed separately on samples within each individual gel.

### Crystallographic analysis of aHE 31 and aHE15

The ancestral reconstructions aHE31 and aHE15 were each purified as fully untagged constructs and purified via heparin affinity chromatography followed by size exclusion chromatography as described above. The proteins were each concentrated to 150 micromolar concentration and mixed in the presence of 10 mM CaCl_2_ with their respective DNA target duplex at 300 micromolar concentration, to achieve a protein:DNA molar ratio of 1:2. Cocrystallization samples were incubated at room temperature for at least 10 minutes prior to setting up crystallization drops. Crystals were grown at room temperature using hanging drop vapor diffusion in a 24-well format.

The sequences of target site DNA duplex oligos for each construct are provided below. For aHE31, crystals were grown in the presence of 28% w/v PEG 3350, 0.2M Ammonium sulfate, 0.1 M HEPES pH 7.5. For aHE15, crystals were grown in the presence of 28% w/v PEG 3350, 150 mM NaCl, 200 mM HEPES pH 7.5. Glycerol (25% v/v) was used as a cryoprotectant for both crystals prior to flash cooling in liquid nitrogen.

aHE31 top strand DNA oligo: 5’ GAACCTTTGGTTATGAGGATCTTC 3’
aHE31 bottom strand DNA oligo: 5’ CGAAGATCCTCATAACCAAAGGTT 3’
aHE15 top strand DNA oligo: 5’ GGTTTACCGATATACACCCCTAAGAG 3’
aHE15 bottom strand DNA oligo: 5’ CCTCTTAGGGGTGTATATCGGTAAAC 3’

Data were collected at the Advanced Light Source synchrotron facility (Berkeley, CA) at beamline 5.0.1 using an x-ray wavelength 0.97Å. The dataset was auto-processed with XDS (data integration and scaling) (57), POINTLESS (space group determination) (58), and AIMLESS (final merging) (59). A signal to noise cutoff of I/sigI >2.0 was applied (corresponding to an ultimate resolution limit of 2.37 Å) and the Phenix crystallography program suite was used (60) for structure determination. Phasing was performed by molecular replacement with Phaser (61) using an AlphaFold-3 generated model of aHE15 as a search model, subjected to AutoBuild, and then the DNA (and all necessary regions of the protein) was manually built and/or rebuilt with iterative rounds of refinement with phenix refine and rebuilding with Coot (62).

Final structures were deposited and validated by the RCSB PDB (RCSB.org) (63) with the PDB identifiers 12FK (aHE31) and 36OF(aHE15). Data and refinement statistics are provided in **Table 1**).

## RESULTS

### 1. Most extant (present-day) LHEs display compromised folding and cleavage activity

We first asked what fraction of monomeric LHEs are capable of being expressed as full-length proteins. Then we then asked how many of the well-expressed proteins display cleavage activity against their predicted DNA target site. We performed these analyses using a dataset of 371 putative LHEs identified via sequence database searches carried out when this project was initiated in 2015. Eight years later, we repeated our database searches, doubling the size of our sequence collection of such sequences. We used this expanded set of sequences for phylogenetic analyses and ancestral reconstruction, as described in the second half of **Results**.

#### Assessment of expression and folding

We were able to successfully subclone 307 of our dataset of 371 candidate LHEs into a yeast surface display vector. Each of those 307 candidate LHE ORFs was assayed for expression on the surface of yeast using flow cytometry to determine the presence or absence of full-length displayed proteins (**Figure 1a**). The folding of protein chains in the eukaryotic endoplasmic reticulum (even those that are not produced via ER transit in their normal physiological context) is subject to rigorous quality control, with misfolded and unstable proteins usually subjected to proteasomal degradation (64); therefore the successful expression of a full-length surface-displayed protein is generally consistent with a properly folded protein chain (65).

The observed expression behavior of the 307 LHEs was binned into three broad overlapping categories (**Figure 1b**): full-length surface expression (staining observed for both N- and C-terminal tags), expression of stalled or truncated constructs (staining observed for only the N-terminal tag), or no surface expression (no staining from either tag). A plot of quantified values for full-length C-terminal staining (**Figure 1c**) reveals a wide range of surface expression signal, with the strongest expressers staining more than two orders of magnitude higher than the weakest. We found that 119 clones (39%) displayed strong signal from both tags, indicating expression of full-length protein on the yeast surface. 142 clones (46%) displayed significantly reduced signal from the C-terminal Myc tag, and the remaining 46 clones (15%) showed no measurable signal above background signal from either the N- or the C-terminal tag.

#### Assessment of cleavage activity

The next question we asked was, for those LHEs that displayed measurable expression on the surface of yeast, what fraction displayed sequence-specific DNA cleavage activity. For that purpose, a total of 188 strongly expressed candidate LHEs (**Figures 1c** and **2c** and **Supplementary Figure S1**) were assayed. To predict a candidate DNA target site for each enzyme, we relied on a well-described aspect of their behavior: because the mobile element inserts into the host gene at the site of cleavage (66), the target sequence can be reconstructed by fusing the host gene sequences that flank the inserted mobile element (i.e. the target sequence corresponds to the intron insertion site in each uninterrupted host gene). The DNA duplexes used in our cleavage assays were all 58 basepairs in length and were centered around the intron insertion site in each corresponding uninterrupted host gene.

We carried out our analysis of cleavage activity using two complementary DNA cleavage assays. In the first approach (**Figure 2ab**), DNA cleavage was measured directly on the surface of yeast using a flow cytometric cleavage assay where a fluorescently labeled, linear DNA target substrate was tethered to the N-terminal HA tag of the displayed enzyme (37). The second approach corresponded to a more traditional “non-tethered” assay in solution, using the same linear DNA target substrates. For this non-tethered assay, the surface-expressed homing endonucleases were released from the yeast surface via treatment with dithiothreitol, and the corresponding DNA target substrate was then added to the supernatant for an *in vitro* cleavage digest reaction. The digest products were visualized via polyacrylamide gene electrophoresis (**Supplementary Figure S2a**).

By employing both of these assays, we could identify active enzymes that might have been missed using only one approach or the other. Those that bind their DNA targets with particularly low affinity may only display cleavage activity when their substrate is physically tethered in proximity to the surface-displayed enzyme. Conversely, those that bind DNA with exceptionally high affinity (and that often do not readily release cleaved DNA products) may only display cleavage activity in a more traditional digest, where bound DNA is physically dissociated from the protein prior to electrophoretic analysis. Altogether, a total of 29 LHEs displayed measurable cleavage activity against their predicted targets. Twenty-four cleaved their DNA targets in both assays, while three (HE07, Gen9-156, and Gen9-171) displayed cleavage activity only in the tethered assay on the yeast surface, and the final two (Gen9-022 and Gen9-051) displayed cleavage activity only in non-tethered *in vitro* cleavage assays.

The DNA target region for each active enzyme was then subcloned into a plasmid backbone, and each plasmid substrate was subjected to a follow-up *in vitro* digest with its corresponding enzyme. Run-off Sanger sequencing of the linearized plasmid product was used to determine the precise location of cleavage on each strand of the DNA and thereby identify the center of each target site and the 4-base overhangs generated by each endonuclease. In the end, we were able to determine the exact center of the target and corresponding sites of DNA strand cleavage for 19 of the 29 active enzymes (**Supplementary Figure S2b**). For the remaining 10 enzymes, we could not obtain sufficient amounts of digested plasmid and/or clear run-off sequencing data, so their targets are listed in **Supplementary Figure S2c** as the full 58 bp DNA substrate sequence (with the site of cleavage expected to occur somewhere within that target region). Raw run-off sequencing data is provided in the **Deposited Raw Data.**

The results of these experiments indicated that (1) as many as 90% of the LHEs in our analysis displayed significantly compromised folding, expression, and/or cleavage activity (presumed to mostly reflect the accumulation of debilitating mutations, although in some cases an error in the assembly of the correct predicted target region sequence would also cause a lack of observed activity), and (2) the remaining 10% of LHEs are expressed well enough to exhibit site-specific DNA cleavage activity. We generated a multiple sequence alignment and phylogenetic tree from the collection of 371 LHEs (**Figure 3a**). The newly identified, active LHEs add siginificant evolutionary diversity to the actuve LHE repertoire: the new active LHEs (red) are distributed around the tree, whereas the previously characterized enzymes (black), which were all identified from a search for close homologues to I-OnuI (37), are found in a fairly narrow subclade of the tree.

### 2. Ancestral reconstruction experiments generate active constructs with enhanced stability and the potential for novel DNA recognition specificity

#### Expansion of the LHE sequence collection, phylogenetic analyses and ancestral reconstructions

We next asked two further questions. First, given that most modern HEs appear to harbor mutations that disrupt folding, stability and/or activity, can well-established methods for generating ancestral reconstructions produce active enzymes? Second, if ancestral reconstructions do in fact yield active enzymes, might some of them reveal additional details of how DNA recognition specificity changes during their evolutionary diversification? To explore these questions, we decided to focus on two subfamilies of related enzymes, one containing I-PnoMI and two newly validated active enzymes (Gen9-348, and HE08, **Figure 3b**), and the other containing the well-studied I-OnuI enzyme (67) and its close homolog I-GpeMI (**Figure 3c**).

Before choosing LHE sequences to use in these ancestral reconstructions, we repeated the bioinformatic search first performed in 2015 in order to obtain a denser sampling of extant sequences. Our 2023 search yielded over 300 additional LHE sequences, so that our newer dataset comprised 622 LHE homologues (**Supplementary Figure S3**). Using those sequences, we extracted the sequences of each LADLIDADG domain (two per single chain monomeric LHE) and generated a phylogenetic tree (**Supplementary Figure S4**). Using that domain-based tree, we identified sequences comprising each subfamily of interest along with selected outgroups (see **Methods**). Using all full-length subfamily sequences as input to the BaliPhy suite (54), we performed multiple sequence alignment, estimated phylogenies and generate predicted ancestral reconstructions at selected nodes, with manual adjustment at positions containing gaps.

#### “Shallow” ancestral reconstructions display enhanced expression, increased thermal stability and retention of activity

The first ancestral reconstruction studied in detail, named ***aHE31***, was generated within the clade containing I-PnoMI, Gen9-348, and HE08 (**Figure 3b**). The pairwise amino acid identity between those three extant enzymes ranges from 63 - 73%; the reconstructed aHE31 has 66 - 78% identity to each of them (**Figure 4a** and **Supplementary Figure S5a)**. The DNA target sites for I-PnoMI, Gen9-348, and HE08 differ at 5 of the 22 basepair positions (**Figure 4b**).

**Figure 4.**
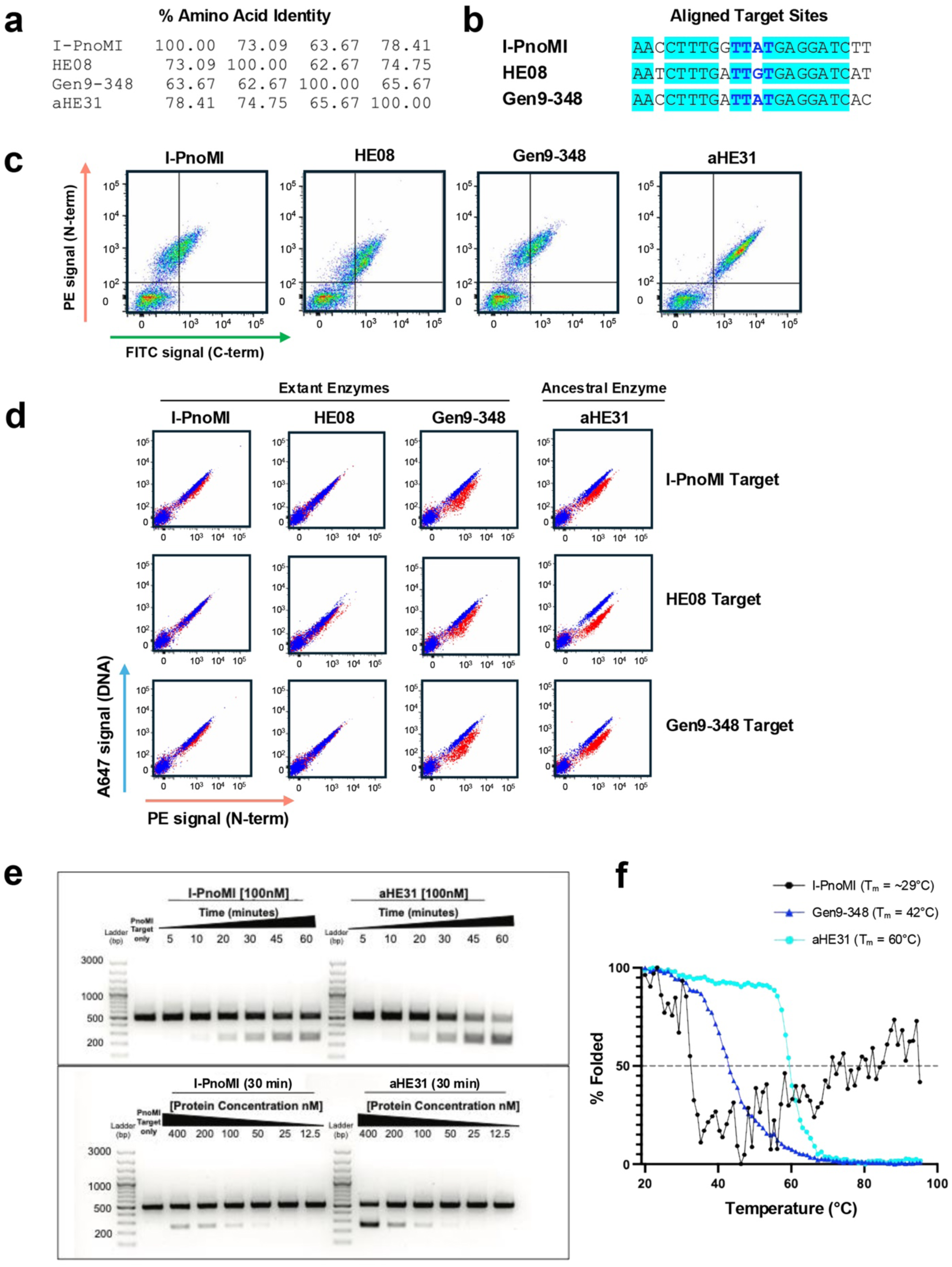
Ancestral reconstruction from closely related enzymes that cleave similar DNA targets. **Panel a:** Table of percent amino acid identity for three modern-day LHEs (I-PnoMI, HE08, Gen9-348) and their reconstructed ancestor aHE31. **Panel b:** Alignment of the verified DNA target sequences of I-PnoMI, HE08, and Gen9-348. The target sequences are identical at 17 of 22 basepair positions (highlighted cyan), with most mismatches occurring on the outer flanks where LHEs characteristically show the most tolerance for sequence changes. There is one mismatch in the “central four” (blue text) of the HE08 target, the region of the target sequence where changes are most likely to disrupt cleavage activity (nonetheless, Gen9-348 can surprisingly accommodate this basepair difference). **Panel c:** Reconstructed ancestor aHE31 and each of the related modern-day enzymes (I-PnoMI, HE08, Gen9-348) were expressed on the surface of yeast and analyzed by flow cytometry for expression (measured by staining the N-terminal HA tag with an anti-HA-biotin/Streptavidin-PE (SAV-PE) conjugation and the C-terminal Myc tag with anti-Myc-FITC, Figure 1a). Full-length expression of the LHE is indicated by the presence of FITC signal. **Panel d:** aHE31 displays robust cleavage of all three extant enzyme target sequences, as measured using the tethered flow cytometric cleavage assay (Figure 2a). **Panel e:** Purified recombinan**t** aHE31 displays a roughly equivalent cleavage rate, and slightly superior cleavage as a function of enzyme concentration, against the I-PnoMI target as compared to the I-PnoMI extant enzyme itself. Cleavage reactions were performed in parallel using equal amounts of a 500 bp DNA substrate containing the I-PnoMI target sequence. Digests were performed at 37°C using 100 nM purified recombinant enzyme with incubation times ranging from 5 - 60 minutes (top two gels) and a range of 12.5 – 400 nM enzyme concentration in digests with an incubation time of 30 minutes (bottom two gels). **Panel f:** Circular dichroism spectroscopy was used to determine the thermal melting temperatures (T_m_) of purified recombinant I-PnoMI, Gen9-348, and aHE31 enzymes. A dashed line marks the temperature at which each protein is 50% unfolded (T_m_): I-PnoMI (∼29°C, estimation only due to instability of the protein in the CD buffer), Gen9-348 (42°C), and aHE31 (60°C).

When all four enzymes are expressed on the surface of yeast, we observed higher expression of the reconstructed ancestor aHE31 than any of its three extant enzyme relatives (**Figure 4c**). We then tested the ability of each of the four enzymes to cleave each of their three closely related DNA targets and observed that aHE31 displayed robust cleavage of each (**Figure 4d**). In contrast, two of the extant enzymes (I-PnoMI and HE08) display slightly lower cleavage activities against a subset of the three targets, while the third (Gen9-348) displayed cleavage activity against all three targets.

To further examine the behavior and activity of aHE31, we generated purified recombinant aHE31 (**Supplementary Figure 5b**) and compared its endonuclease activity to that displayed by recombinant I-PnoM against that enzyme’s DNA target sequence. In cleavage assays conducted over a range of digest times and enzyme concentrations, aHE31 displayed roughly equivalent cleavage rates and yields of product formation as a function of both digest time and decreasing enzyme concentration (**Figure 4e**).

We examined the thermal stability of aHE31 by determining the thermal denaturation temperature and unfolding behavior of aHE31 compared to modern-day enzymes I-PnoMI and Gen9-348 (**Figure 4f**). A thermal denaturation curve was not obtained for HE08 because that protein could not be adequately expressed and purified from bacteria. The purified recombinant I-PnoMI protein was markedly unstable in the CD buffer, and the resulting melting curve barely allowed for the determination of an estimated T_m_ value of ∼29° C. The melting curve for Gen9-348 displayed a much better overall shape with cooperative unfolding and a T_m_ value of 42°C. In contrast, the T_m_ of aHE31 (60° C) was significantly increased relative to I-PnoMI and Gen9-348 (similar increases in thermal folded stability are often observed for ancestral reconstructions (68) and prove valuable for enzyme and protein engineering studies (69)). The crystal structure of aHE31, bound to the I-PnoMI target site, was determined to 2.17 Å resolution (**Table 1** and **Supplementary Figure S5c**) and found to be closely superimposable to the previously determined structure of I-PnoMI (70), as well as more distantly related LHE homologues such as I-OnuI (67).

A second ‘shallow’ ancestral reconstruction (***aHE11)*** was also examined, to assess the reproducibility of the observations described above. Similar to aHE31, the aHE11 construct corresponded to a node located a short step backwards in the phylogenetic tree, near a pair of even more closely related extant enzymes (I-OnuI and I-GpeMI) (**Figure 3c**). Those two modern day LHEs are 86% identical to each other and recognize DNA target sites that differ at only two positions (**Figure 5ab**). aHE11 is 93% identical to each of those enzymes. Like aHE31, aHE11 displays a higher level of full-length expression on the surface of yeast than its descendants (**Figure 5c**), cleaves both the I-OnuI and I-GpeMI targets (**Figure 5d**), and displays well-behaved, cooperative unfolding corresponding to a thermal denaturation temperature (T_m_ = 50°C) that is slightly increased relative to the previously determined denaturation temperature for I-OnuI (T_m_ = 46°C) (29) (**Figure 5e**).

**Figure 5.**
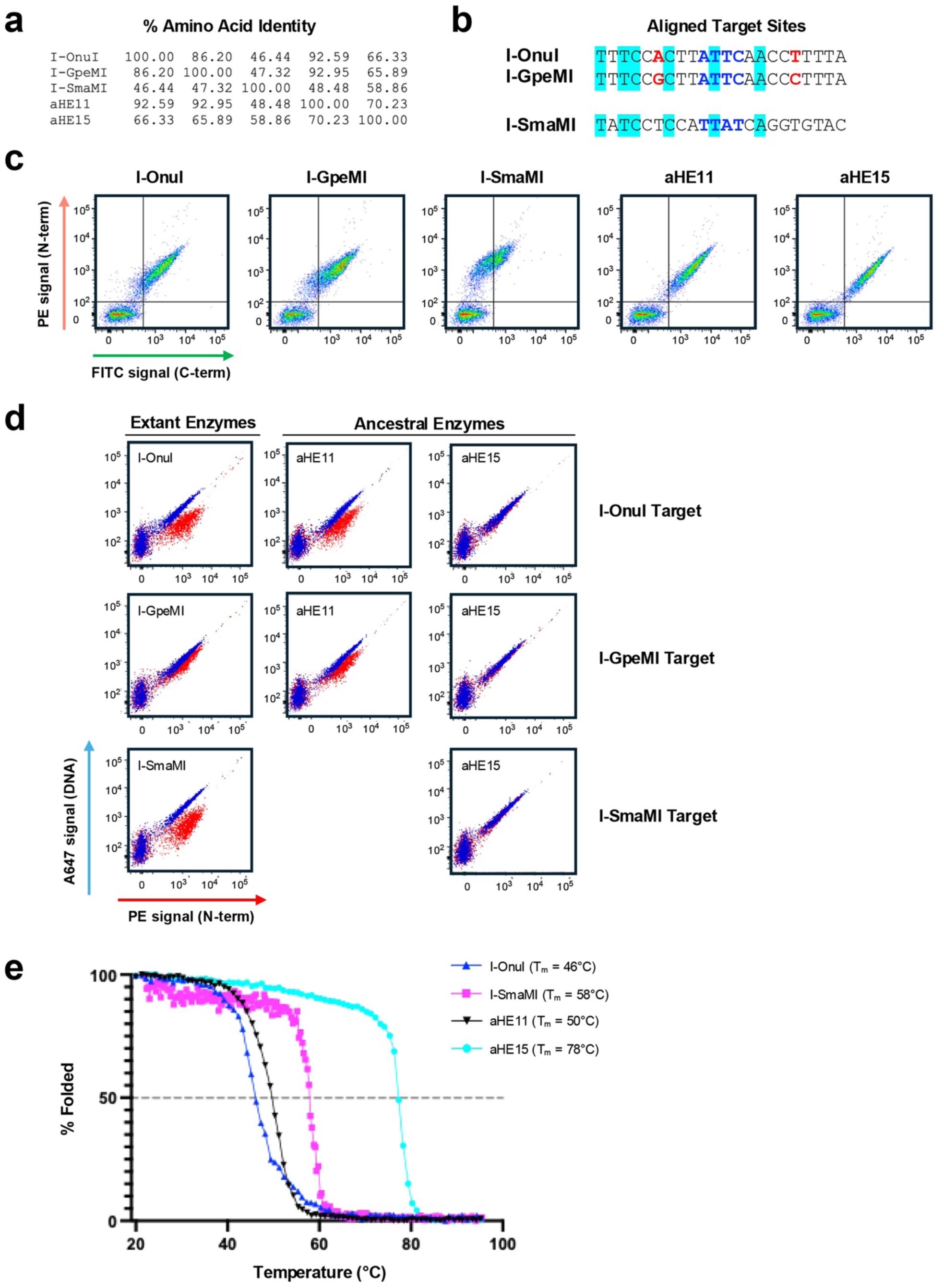
Ancestral reconstructions from a more divergent group of LHEs. **Panel a:** Table of percent amino acid identity of three modern day LHEs (I-OnuI, I-GpeMI, I-SmaMI) and their reconstructed ancestors aHE11 and aHE15. **Panel b:** Alignment of the verified DNA target sequences of I-OnuI, I-GpeMI, and I-SmaMI. The target sequences of I-OnuI and I-GpeMI are identical at 20 of the 22 basepair positions, with 2 mismatches highlighted in red. The DNA target sequence of the more distantly related I-SmaMI matches I-OnuI and I-GpeMI at only 6 of 22 basepair positions (highlighted in cyan). **Panel c:** Reconstructed ancestors aHE11and aHE15 are expressed on the surface of yeast at equal or superior levels as compared to each of their related modern-day enzymes (I-OnuI, I-GpeMI, I-SmaMI). Expression was measured as described in methods and illustrated in Figure 1a. **Panel d:** Shallow ancestral reconstruction aHE11 displays robust cleavage of either of the target sites for its two immediate descendants (I-OnuI and I-GpeMI). In contrast, the deeper ancestral reconstruction aHE15 does not display cleavage of any of the target sites for its more distant ancestors (I-OnuI, I-GpeMI or I-SmaMI). using the tethered flow cytometric cleavage assay (Figure 2a) described in **Methods. Panel e:** Reconstructed ancestors aHE11and aHE15 display increased thermal folding stability relative to their extant modern-day counterparts, as evaluated using circular dichroism spectroscopy (CD) to determine the thermal melting temperatures (T_m_) of purified recombinant enzymes. Melting data was plotted alongside previously collected data for related modern-day LHEs I-OnuI and I-SmaMI (29). A dotted line marks the temperature at which each protein is 50% unfolded (T_m_): I-OnuI (46°C), I-SmaMI (58°C), aHE11 (50°C), and aHE15 (78°C).

**Figure 6.**
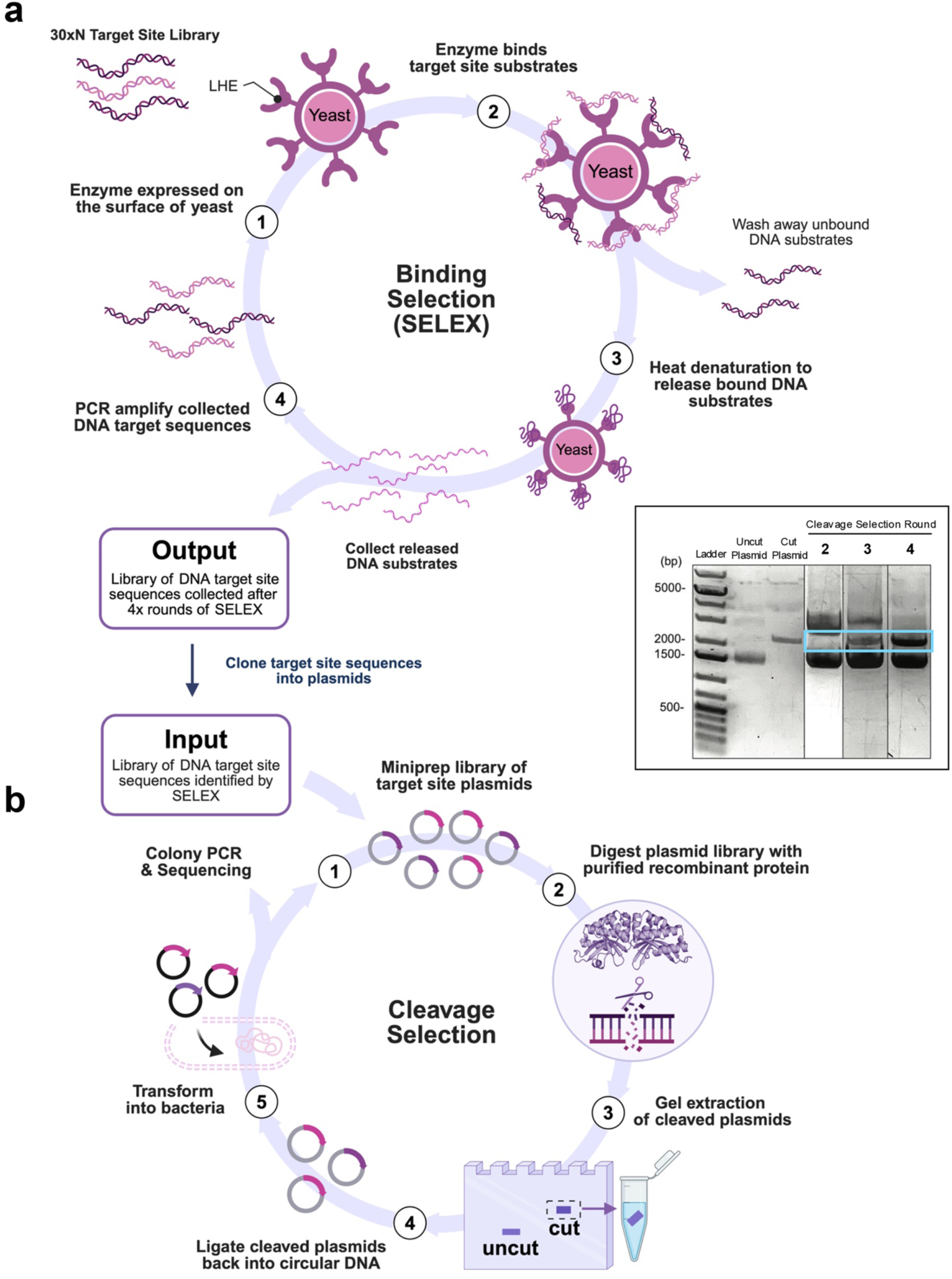
Two-stage selection strategy for determination of a cleavable DNA target sequence for ancestor aHE15. **Panel a:** Schematic of the binding-based SELEX protocol used to isolate a pool of DNA target substrates bound by the aHE15 enzyme. (1) Ancestor aHE15 was expressed on the surface of yeast and incubated with a library of double-stranded DNA target substrates containing a fully randomized 30 bp sequence (30xN) flanked by two primer sequences (**Supplementary Figure S8a**). (2) aHE15 was allowed to bind compatible DNA target substrates and all remaining unbound DNA was washed away. (3) Yeast with substrate-bound aHE15 was heated to denature the enzyme and release any bound DNA substrates. (4) The released DNA target substrates were collected and PCR-amplified to create an enriched pool of DNA target substrates for the next round of SELEX. This process was repeated for a total of 4 rounds, and the resulting pool of DNA target sequences bound by aHE15 was cloned into plasmids and moved on to the cleavage selection stage. **Panel b:** Schematic of the cleavage-based selection protocol used to isolate DNA sequences cleavable by the aHE15 enzyme. (1) A library of 30 bp DNA sequences bound by surface-expressed aHE15 in the SELEX protocol was cloned into plasmids and miniprepped to prepare the substrate for cleavage digests. (2) The library of target site plasmid DNA was digested with purified recombinant aHE15 enzyme. (3) The digest reaction was loaded onto an agarose gel, and any linearized plasmids were isolated by gel extraction. (4) The cleaved/linear target site plasmids were purified and re-ligated back into circular DNA. (5) The circularized plasmids containing cleavable target sequences were transformed into bacteria. A portion of the transformation reaction was plated to allow for colony PCR and sequencing of individual targets, and the remainder was grown overnight in liquid culture to be used for a miniprep to prepare the enriched substrate for the next round of cleavage selection. This process was repeated for a total of four rounds. **Inset:** Image of the plasmid digest reactions from cleavage selection rounds 2 through 4 run out on an agarose gel prior to extraction of any linearized product. A blue box indicates the expected location of linearized plasmid, which was guided by a PvuI-linearized control sample in the adjacent lane as well as an additional reference lane containing uncut/supercoiled plasmid. Cleaved plasmid was only visible after round 3, and the gel for Round 1 is not shown. This image was created from separate gels run on different days.

### A “deeper” ancestral reconstruction displays robust expression and folding, broad DNA binding specificity, and a novel DNA cleavage specificity

A third and final ancestral reconstruction, aHE15, resides in the same clade as aHE11 (**Figure 3c**), but is a deeper (older) ancestor that connects I-OnuI and I-GpeMI to one of the outgroup sequences used for this clade, an active LHE named I-SmaMI (71). I-SmaMI has a more highly diverged protein sequence relative to I-OnuI and I-GpeMI and significantly different DNA target specificity (**Figure 5ab** and **Supplementary Figure 6a**). I-OnuI and I-GpeMI do not cleave the target site of I-SmaMI, and vice-versa.

aHE15 displayed visibly higher expression on the surface of yeast as compared to I-OnuI and I-GpeMI, similar expression to its shallower ancestral neighbor, aHE11, and significantly higher expression than I-SmaMI (**Figure 5c**). Unlike the “shallow” ancestral reconstructions described above, surface-expressed aHE15 did not demonstrate measurable cleavage of any of the DNA targets for I-OnuI, I-GpeMI, or I-SmaMI (**Figure 5d**). We therefore examined the ability of aHE15 to bind any of those three targets, despite its lack of cleavage activity. We observed that aHE15 bound to all three targets when the DNA substrate was provided at concentrations of 25nM or higher, with binding to the I-SmaMI target being the strongest (**Supplementary Figure S7**).

We expressed and purified recombinant aHE15 (**Supplementary Figure S6b**) and determined the thermal denaturation behavior and corresponding T_m_ values for that construct, two of its modern-day extant relatives, and the shallower ancestor aHE11 (**Figure 5e**). The thermal stability of aHE15 was found to be significantly higher (T_m_ = 78°C) than I-OnuI (T_m_ = 46°C), aHE11 (T_m_ = 50°C), and I-SmaMI (T_m_ = 58°C).

Given that ancestor aHE15 did not display cleavage activity against the DNA targets of its three nearest characterized ‘descendants’ (even though it displayed binding to each), we hypothesized that the DNA cleavage specificity for this deeper ancestral reconstruction (if it was indeed active) might differ significantly from those extant enzymes. We therefore employed an unbiased two-stage selection approach (see **Methods** and **Figure 6**) to determine the cleavage activity and specificity of aHE15. That strategy consisted of: (1) four rounds of SELEX DNA target <u>binding</u> selection against a fully randomized 30xN bp DNA target substrate library (**Supplementary Figure S8**), followed by (2) four rounds of a DNA target <u>cleavage</u> selection using a plasmid library containing the target sequences previously isolated via the SELEX binding protocol. The concept of this screen was to (1) reduce the initial complexity of the 4^30^ DNA library (spanning roughly 1.15 × 10^18^, or 2 micromoles, of unique DNA target sequences) to only those sequences that could be bound by the aHE15 enzyme with roughly low micromolar affinities under moderate stringency (300 mM KCl, pH 7.4.), and then (2) pass on those bound DNA target candidates to a secondary screen to identify DNA targets that the aHE15 enzyme could also cleave.

Cleavable DNA targets were monitored after each round of the cleavage selection by sequencing recovered DNA target sites and looking for enrichment of cleavable sequences. After four rounds of cleavage selection, there were 3 cleavable target sequences that appeared at least twice, with the two most highly enriched sequences present 4 times (“4xSeq”) and 17 times (“17xSeq”) in 29 sequence reads (**Supplementary Figure S9**). Purified recombinant aHE15 was used to cleave the target site plasmids for each of these two enriched sequences, and run-off sequencing was performed to determine the precise position of cleavage within each target sequence and then to infer the enzyme’s 22bp target site within the 30bp selected sequence (**Supplementary Figures S10** and **S11**). Alignment of the centered 22 bp 4xSeq and 17xSeq sequences with one another revealed that they are identical at 14 of the 22 positions, demonstrating that our cleavage screen was converging on a pool of similar targets (**Supplementary Figure S12**). Further, both sequences could be aligned with the target sites of I-OnuI and I-SmaI (**Supplementary Figure S12**).

Next, the 22 bp 4xSeq and 17xSeq targets were generated as short linear fluorescently-labeled DNA substrates and used in cleavage digests to determine which sequence was cleaved most efficiently. The more highly enriched target site sequence (17xSeq) was confirmed to be the superior substrate. In the same experiments, we also observed that purified recombinant aHE15 enzyme, at high enzyme concentration and over an extended digest time, displayed very low cleavage activity against the I-OnuI DNA target sequence (that was not previously observed on the surface of yeast in a tethered cleavage assay), while exhibiting no measurable cleavage of the I-SmaMI target sequence (**Supplementary Figure S12**).

### Determination of the DNA cleavage specificity profile of the aHE15 reconstructed ancestor

Having identified a cleavable DNA substrate sequence for the aHE15 enzyme, we next asked (1) whether the corresponding specificity profile displayed by the enzyme (i.e. its sensitivity towards single basepair substitutions at each position in its target) resembles those of previously examined modern-day LHEs, and (2) whether the selected target sequence represents an optimal target site for the enzyme (or if one or more single base substitutions would further increase the target’s cleavability). We therefore performed a “one-off” cleavage specificity assay, using the 17xSeq target sequence as the starting point and testing 66 different target substrates corresponding to the systematic incorporation of each of three alternative bases at each of the target’s 22 individual positions.

The resulting specificity profile for aHE15 (**Figure 7a**) qualitatively resembles those previously established for both I-OnuI and I-SmaMI (37,67), including strong sensitivity towards individual base pair substitutions (resulting in significantly reduced cleavage) across the central four basepairs of the target and many positions extending outwards, contrasted by reduced fidelity at the outer-most two or three base pair positions in each DNA half-site. At least four individual basepair substitutions (-4C to A, +4A to T , +4A to C, and +10T to A) increased cleavage efficiency relative to the original 17xSeq target sequence. We therefore tested an additional seven target sequences that included all possible combinations of each of the single basepair substitutions (**Figure 7b** and **Supplementary Figure S13ab**). That secondary set of assays indicated that the most efficiently cleaved DNA target sequence for aHE15 corresponded to 5’ TTTACCGATATACACCCCTAAG 3’. That target site shares 54% identity with the target of I-OnuI (12 out of 22 base pairs) and 27% identity with the target of I-SmaMI (6 out of 22 base pairs) (**Figure 7c**).

**Figure 7.**
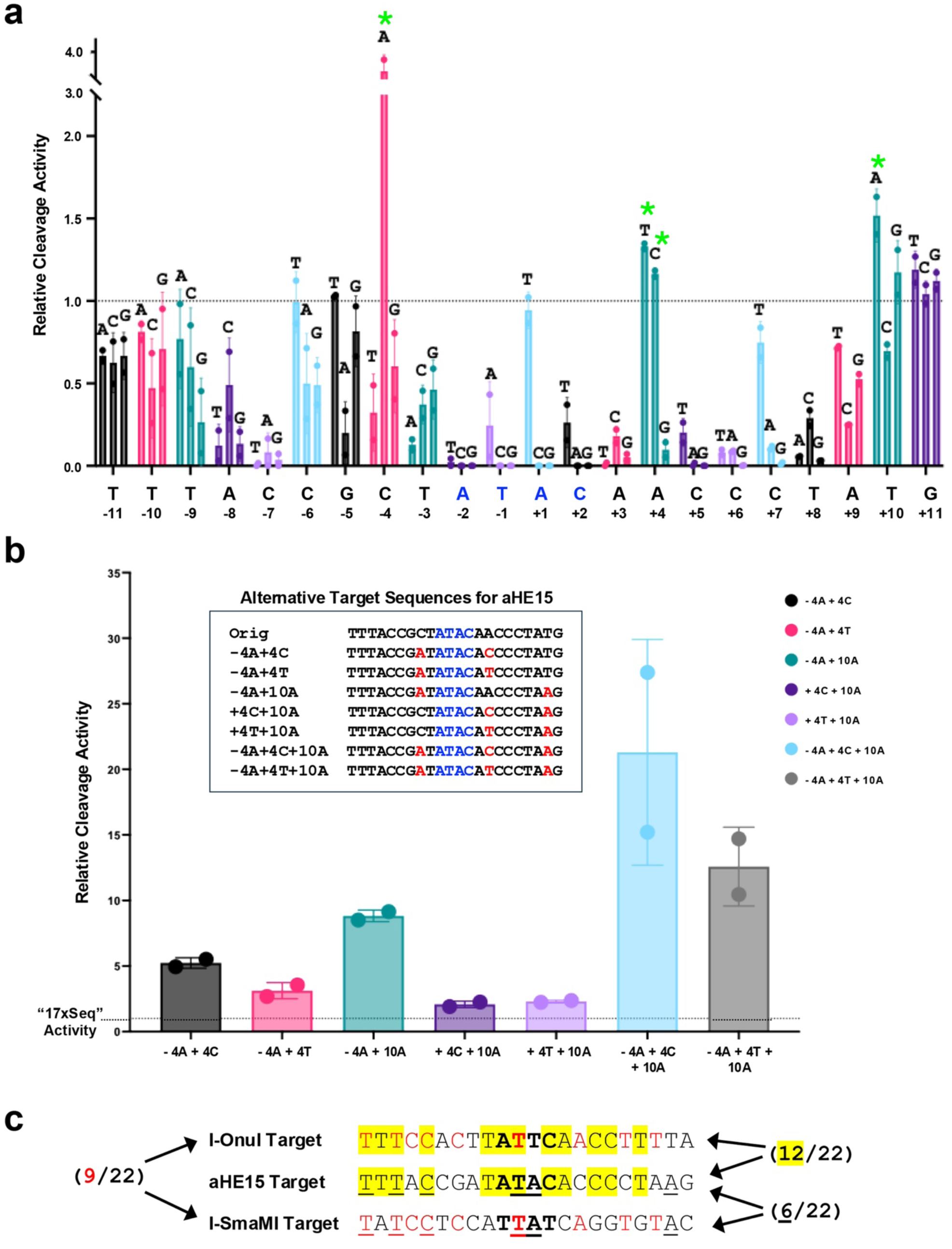
One-off specificity profile for aHE15. **Panel a:** The DNA cleavage specificity profile for aHE15 resembles previously described specificity profiles for modern-day LHEs. Recombinant purified aHE15 enzyme was used to cleave a set of 66 different DNA target substrates, each with a single basepair change away from the most enriched (17xSeq) target sequence from our screen. The sequence of 17xSeq is shown on the x-axis along with numbered positions in the target. The central four bases of the target sequence are colored blue. Each bar represents the relative cleavage (compared to cleavage of the original 17xSeq sequence) of each one-off DNA substrate, with the alternative base indicated above each bar. The dotted horizontal line designates a relative cleavage value of 1.0, or the level at which the aHE15 enzyme cuts the original 17xSeq target sequence. Four alternative one-off targets cleaved significantly better than 17xSeq: -4C, +4C, +4T, and +10A (green asterisks). **Panel b**: Relative cleavage activity displayed by aHE15 against seven additional target sequences (listed in the **Inset**), that were generated by incorporating all combinations of the four improved one-off substitutions, led to the identification of the final ‘optimal’ DNA target that was used for structural analyses (Figures 8 **and 9**). Recombinant aHE15 enzyme was used to digest each of the alternative combinatorial target sequences, and the intensity of cleavage bands on a gel (**Supplementary Figure S13**) was quantified and normalized to that of the original 17xSeq target sequence (dotted line). **Panel c**: Alignment of the final ‘optimal’ DNA target for aHE15 to the DNA targets previously identified for extant enzymes I-OnuI and I-SmaMI. The central four bases of each target are shown in bold text. Nine matches between the I-OnuI and I-SmaMI targets are colored with red text, 12 matches between the aHE15 and I-OnuI targets are highlighted yellow, and 6 matches between the aHE15 and I-SmaMI targets are underlined.

### Structural features of DNA recognition and its divergence

Having determined a DNA target sequence that was near-optimal for the reconstructed ancestral LHE construct aHE15, we co-crystallized the protein bound to its DNA target site and solved the structure to 2.37 Å resolution. The final refined model (**Figure 8a**) revealed a single copy of the aHE15 construct bound in an unambiguous orientation to a single copy of its DNA target site, with the central four basepairs and corresponding scissile phosphate groups on each strand centered within the enzyme’s endonuclease actives sites as previously observed in the DNA-bound structures of its two previously visualized “descendants”, I-OnuI and I-SmaMI (37,67).

**Figure 8.**
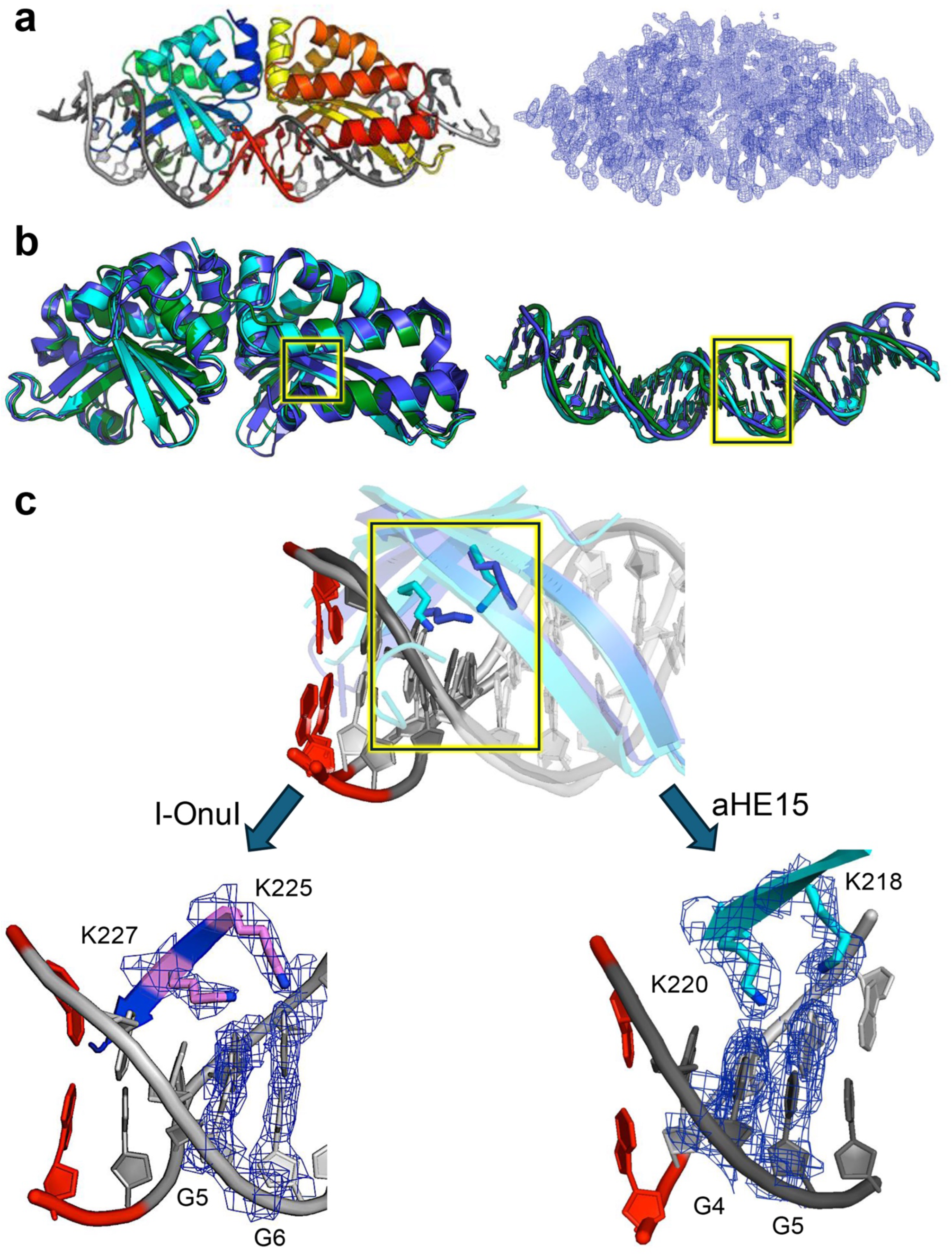
Crystal structure of aHE15 bound to its DNA target. ***Panel a*:** Ribbon diagram of aHE15 (colored in a rainbow scheme denoting the progression of the protein chain from the N-terminus (blue) to the C-terminus (red) and corresponding 2Fo-Fc composite omit map of the protein-DNA complex, contoured at 2σ. The central four bases of the bound DNA target are colored red. ***Panel b***: Superposition of the protein (left) and DNA (right) components of the I-OnuI (blue), I-SmaMI (forest green) and aHE15 (cyan) protein-DNA complexes. ***Panel c***: Superposition of the DNA-contacting beta-strand spanning equivalent lysine residues from I-OnuI (K225 and K227) and aHE15 (K218 and K220), demonstrating different rotameric conformations and corresponding contacts to different DNA basepairs within those enzymes’ respective targets. The corresponding regions of protein and DNA target, and corresponding 2Fo-Fc electron density for each, are shown below the superpositions. The highlighted boxes in panel b correspond to the regions shown in Panel c.

Whereas the conformation of the aHE15 protein was closely superimposable to DNA-bound structures of both I-OnuI and I-SmaMI (**Figure 8b**; pairwise superposition RMSD values against either of those enzymes, calculated for all comparable α-carbons, were both approximately 0.9 Å), the corresponding superimposed DNA targets displayed more significant local differences in their conformations (pairwise superposition RMSD values both approximately 2 Å), with much of the difference in DNA conformation attributable to differences in major groove dimensions along the length of the respective DNA targets.

The identity and details of individual DNA contacts formed by I-OnuI, I-SmaMI and their common reconstructed ancestor aHE15 display a mixture of conservation and divergence from one another (**Figure 8c** and **Figure 9**). All three enzymes rely on approximately ten residues, largely located at comparable positions in their aligned structures, to form direct side chain contacts with individual DNA bases. Those contacts are augmented by water-mediated contacts between additional protein side chains and individual DNA bases and further sequence-nonspecific contacts to the DNA backbone, collectively involving roughly an additional 20 residues in DNA contacts and binding. Those roughly 30 DNA-contacting residues in each structure are in contact with an additional 20 neighboring residues that provide structural context that reinforces the overall shape and stability of the DNA-contacting surface of the protein.

**Figure 9.**
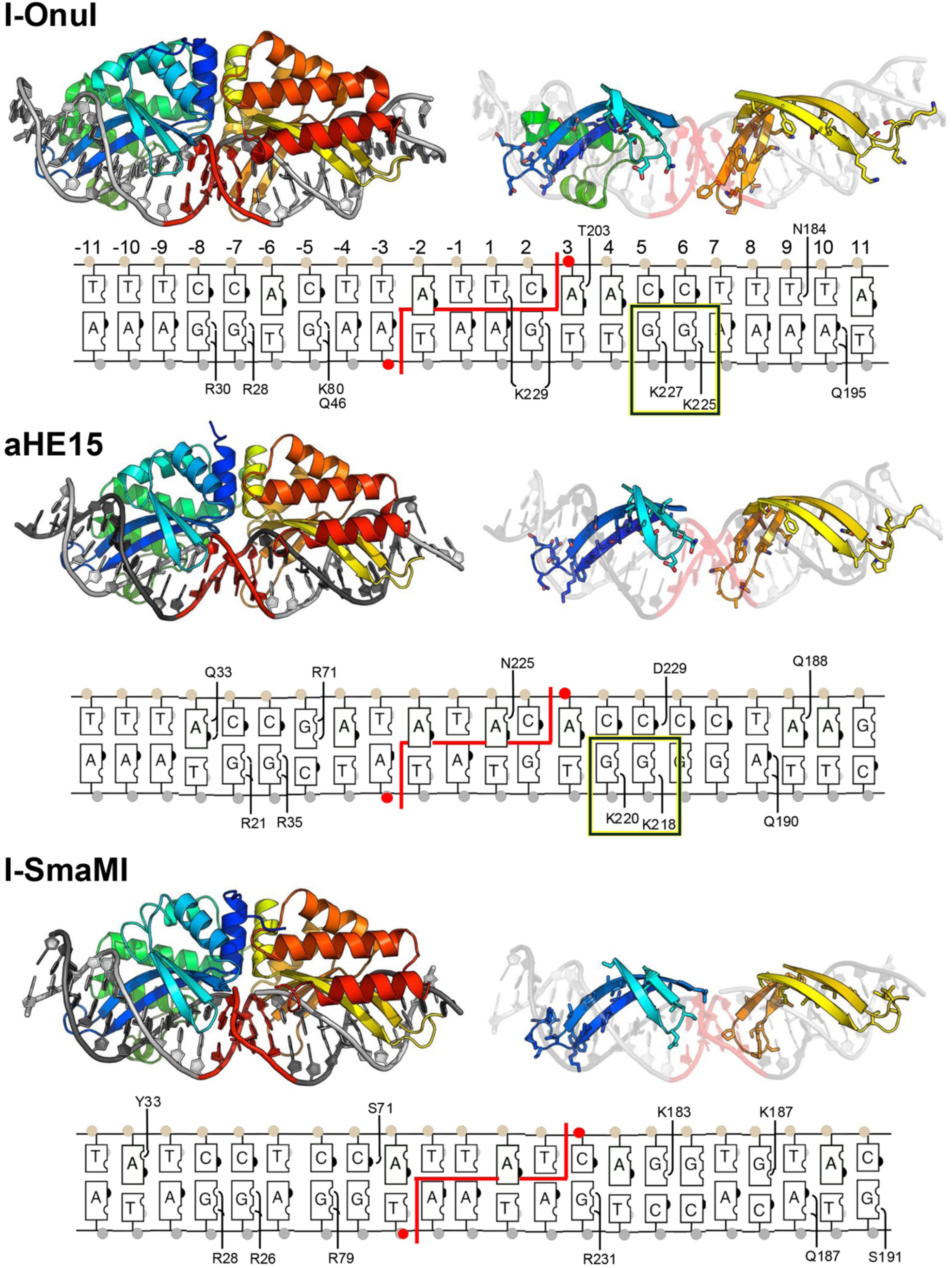
Protein-DNA contacts. Cartoon renderings of the intact protein-DNA complex and of the DNA-contacting β-sheets of each enzyme (I-OnuI, I-SmaMI and aHE15) are shown above corresponding contact schematics for each enzyme. For clarity, only direct contacts between protein side chains and corresponding DNA bases are indicated. Those contacts are joined by an additional set of water-mediated contacts between additional protein side chains and DNA bases, and further contacts to the DNA backbone, ultimately involving approximately 30 residues of each protein (approximately 10% of the total protein residues in each structure). The highlighted boxes also correspond to the regions and contacts that are shown in Figure 8c.

The identity of some, but not all, of the individual residues that are in contact with DNA bases in each structure are conserved -- for example, lysines K225 and K227 in I-OnuI versus lysines K218 and K220 in aHE15 (**Figure 8c**) or arginines R28 and R30 in I-OnuI versus arginines R26 and R28 in I-SmaMI. Although conserved, the sidechains of those residues often display differing rotameric conformations that place them in direct contact with chemically identical, but differently located base pairs within each enzyme’s target site. For example, K225 and K227 in I-OnuI form contacts to guanine bases at positions +5 and +6 in that enzyme’s target, whereas comparable lysine residues K218 and K220 in aHE15 make similar contacts to guanine bases at positions +4 and +5 in its target (**Figure 8c** and **Figure 9**).

Conversely, whereas R28 and R30 in I-OnuI (and R26 and R28 in I-SmaMI) make similar direct contacts to guanine bases located at positions -7 and -8 in their respective targets, aHE15 utilizes arginines at two nearby but different positions (R21 and R35) to form similar direct contacts to guanine bases at positions -6 and -7 in its own DNA target. Such indications of structural malleability in contacts between protein side chains and DNA bases have been previously observed for various related LHEs (31,37,72).

Our observations of malleability in the LHE-DNA target contact map between a present-day LHE and its predicted ancestor reinforce previous studies that found similar divergence in protein-DNA contacts between naturally evolved LHE homologues (37) and between different laboratory-engineered LHE-DNA complexes (31).

## DISCUSSION

### Homing endonuclease diversity and evolutionary degradation

Most mobile genetic elements (MGEs) provide little or no benefit to their host and are under minimal selection pressure to maintain their form and function, thereby leading to the accumulation of debilitating mutations and eventual inactivation (73). For example, LINE-1 retrotransposons represent 17% of the human genome, but fewer than 1% are active, due to truncations, internal rearrangements, and point mutations (74). LINE-1s have even become totally ‘extinct’ in some mammalian lineages (75). Similarly, the human genome harbors many fossil DNA transposon sequences, but none are now active (having become extinct over the past ∼40 to 50 million years (2).

The presence, action, and evolution of MGEs extends to (and likely originated in) the microbial universe; the life cycle and distribution of their activities and dysfunctions have been examined in that context as well. Early studies of LHE life cycles, beginning with an analysis of the yeast I-SceI homing endonuclease lineage (and their corresponding mobilized introns) indicated that they undergo an evolutionary life cycle of invasion, mutational degradation, loss from the genome, and subsequent re-invasion by a related element, with an estimated periodicity of approximately 2 million years (32). A similar study of the I-OnuI homing endonuclease examined a group of closely related homologues from *ascomycete* fungi and concluded that over half appeared to harbor inactivating genetic truncations and mutations (76). Yet another analysis of a small collection of mobile introns that reside in rDNA host genes, and that are mobilized by members of the “His-Cys box” homing endonuclease family, arrived at similar numbers and conclusions (77).

Those prior studies of LHEs relied on approaches in which a relatively small number of homologues were characterized, in each case closely related to least one highly active homing endonuclease. To more broadly examine how frequently such elements retain both form and function, we applied a strategy in which a much larger, unbiased collection of LHEs was identified based on simple criteria that (1) they are encoded within group I introns (that often act as a surrounding vehicle for homing endonuclease reading frames), and (2) that they appear to be members of the single-chain (monomeric) LAGLIDADG homing endonuclease (LHE) family. We cloned 307 candidate enzymes, assayed each for protein expression and folding, and further assayed the well-expressed subset for DNA cleavage activity against their predicted target sequences.

We found that roughly 90% of these 307 homing endonuclease genes (HEGs) display significant deficiencies in their expression, folding, stability, and/or DNA cleavage activity. Furthermore, while expression of full-length protein is obviously a necessary prerequisite for cleavage activity, a significant fraction of the HEGs that displayed robust expression and folding nonetheless displayed little or no endonuclease function (**Figure 2c**). The 29 newly identified active endonucleases are distributed throughout the LHE phylogenetic tree, in contrast to previously identified active LHEs which are found in a more restricted subclade (**Figure 3a** and **Supplementary Figure S3**). Each new active enzyme cleaves a DNA target corresponding to the mobile intron insertion site within its host gene. This expanded range of DNA target sequences might provide an excellent starting point for engineering enzymes with new, useful specificities.

### Ancestral reconstructions yield LHEs with robust expression and cleavage activity

Ancestral sequence reconstruction (ASR) methods infer the most probable sequences of extinct ancestral molecules, by combining alignments of extant homologues with phylogenetic models of sequence evolution (78). Modern ASR methods typically use likelihood- or Bayesian-based approaches to estimate the most likely ancestral states at internal nodes of an evolutionary tree, generating hypotheses about historical protein or nucleic acid sequences that can subsequently be evaluated computationally or experimentally. Reconstruction accuracy and confidence depend on dense taxonomic sampling, high-quality multiple sequence alignments, accurate phylogenetic models, and the existence of sufficiently conserved homologous families. ASR therefore performs best when proteins being used for that purpose have evolved under functional constraint (purifying selection) because phylogenetic signal is preserved and historical states can thereby be reliably inferred (79).

In this context, successful ancestral reconstruction of mobile elements might have been expected to prove difficult. Unlike many protein families that evolve under sustained functional constraint, selfish mobile elements experience episodic bursts of positive selection during invasion and spread, followed by prolonged periods of neutral evolution after their fixation. As a consequence, many mobile elements accumulate substitutions, indels and truncations that reduce expression, stability or catalytic activity, and ultimately become inactive prior to elimination from the host genome (17). These mutations might erode the phylogenetic signal required for accurate ancestral inference.

Despite such reservations, we noted that ancestral sequence reconstructions have previously been applied to mobile genetic elements to predict and experimentally evaluate ancestral states using collections of extant, often degraded, sequences (for example, (80–82). Conceptually, many ancestral nodes in a mobile element phylogeny represent transposition events that led to new element insertions, and thus by definition represent active enzymes (while other nodes may represent copies derived by speciation events, i.e. divergence of host genomes rather than of active mobile element copies).

Ancestral reconstructions therefore (if accurate) should be enriched for active enzymes when compared to extant sequences. Collectively, those studies illustrate the potential of ancestral reconstructions of such elements: resurrection of ancestral molecules can reveal biochemical activities no longer represented among extant sequences, but reconstruction fidelity ultimately depends on retaining sufficient phylogenetic signal despite subsequent sequence degeneration.

In this study, we apply ancestral reconstruction to the LHE family of mobile elements, performing detailed functional characterization of one ‘deep’ and two ‘shallow’ ancestral sequences. All three enzymes express and fold well, and exhibit robust DNA target site cleavage activity. One of the three ancestral reconstructions targets anovel DNA sequence not cleaved by its descendent enzymes. Theses successful ‘resurrection’ indicates that sufficient evolutionary information remains to recover coherent ancestral states, indicating that ancestral reconstruction can provide an effective strategy for recovering latent functional diversity from degraded populations of selfish genetic elements, offering access to evolutionary intermediates and activities that have been lost during the natural life cycle of mobile endonucleases.

However, it is worth noting that while the reconstruction effort leading to the “deeper” aHE15 construct was successful in generating a functional enzyme that displays a DNA recognition and cleavage specificity profile that is nearly indistinguishable from modern-day extant LHEs, that its enzymatic power (as measured by its rate of DNA cleavage) as compared to its “descendants” is slightly lower than its modern-day relatives, both on the surface of yeast and in assays using purified recombinant protein. One possibility is that the features we would consider to be optimal for engineered enzymes may not be truly optimal features for the evolutionary success of a mobile element. Maximal expression and cleavage activity may result in mobile elements that are ‘too efficient’, harming their hosts through excessive insertions. Lower levels of activity may be more evolutionarily successful, allowing the element to create new copies at appreciable levels without population-level damage to the host.

Alternatively, technical considerations may mean we have not reconstructed a true ancestor. While aHE15 clearly possesses the necessary components of a LAGLIDADG endonuclease active site, and displays DNA binding affinity and specificity that also approximates the behavior of current day enzymes, it may be that subtle features of structure and dynamics that are thought to be critical for optimal enzymatic function are not fully recapitulated in the computations leading to the final version of such a reconstructed enzyme. Each ancestral sequence we tested is only one of many possible solutions to the true sequence at that node of the phylogeny. While at each protein position, the algorithm chooses the most likely amino acid, very few positions are predicted unambiguously, and the combinatorial uncertainty over the entire protein means that the single full-length sequence prediction we tested is highly unlikely to exactly match the true ancestral sequence. Furthermore, the deeper the node being predicted, the greater the uncertainty. It is also possible that we have not identified an optimal DNA target sequence for the enzyme, and that deeper exploration of DNA sequence space around the current ‘best’ substrate might yield further improvements in endonuclease function.

Finally, this work lays the groundwork for future uses of ancestral reconstruction to understand LHE biology and to aid in LHE engineering. For example, one could reconstruct stepwise series of ancestral nodes in order to examine how these enzymes evolve to change their DNA targeting specificity, and how they acquire the ability to invade novel host genomes. Understanding the molecular patterns that govern enzyme-target recognition could help engineer enzymes with novel desired target sites. While our current work reconstructs ancestors of active descendant enzymes, we also anticipate that ancestral reconstructions would also succeed in clades that contain only ‘dead’ enzymes, potentially increasing the sequence space of active enzymes even further. Furthermore, ancestral reconstructions of such elements can yield enzymes with new DNA targeting specificities, as well as highly expressed, thermostabilized predecessors of compromised modern-day enzymes. Given the size of the current collection of LHEs represented in this study (both modern-day enzymes at the tips of the branches of the phylogenetic tree, along with the underlying nodes that connect every branch point in that tree), it seems possible that this system, like many others of similar diversity and number, may offer considerable potential for the future computational design of a wide range of compact gene-targeting proteins with similarly diverse DNA sequence targets, as well as the potential to readily re-activate and characterize the specificity of many hundreds of such enzymes.

## Supporting information

Supplementary sequence information

## Acknowledgements

We thank Professor Harmit Malik at the Fred Hutchinson Cancer Center for advice and support, and members of his group, as well as those of the Brett Kaiser research group at Seattle University and the Stoddard group at Fred Hutchinson Cancer Center) for additional advice and discussion, particularly Alex J. Kaiser for assistance with subcloning ancestral reconstructions and assistance with protein expression and purification.

## Author contributions

JCY and ARL contributed equally. DRE identified the initial collection of LHE sequences. ARL carried out the cloning and initial characterization of the expression and cleavage activity displayed by that set of LHE sequences, and identified additional LHE sequences for further analysis. JMY curated the corresponding LHE sequence collection and carried out sequence analyses, phylogenetic tree construction and generation of sequences of ancestral reconstructions. JCY carried out biochemical and structural analyses of ancestral LHEs. MS, LAD and BLS assisted with protein purification, biochemical characterization and structural analyses. BLS supervised the project and obtained funding support. All authors contributed to data analysis and writing the manuscript.

## Funding

Funding support was provided for this project by bluebird bio inc., the Fred Hutchinson Cancer Center, and NIGMS grant R35 GM148166 to BLS, NIGMS grant R01 GM074108 to JMY (PI: Harmit S. Malik), NCI T32 CA080416 to JCY, and the NSERC Discover Grant RGPIN-2015-04800 and RGPIN-2022-05459 to DRE. Crystallographic data was collected at the Fred Hutchinson Cancer Center home source (supported in part by a shared instrumentation grant (NIH S10OD028581) and the Advanced Light Source (ALS) synchrotron facility. The ALS-ENABLE beamlines are supported in part by the National Institutes of Health, National Institute of General Medical Sciences, grant P30 GM124169-01. The Advanced Light Source is a Department of Energy Office of Science User Facility under Contract No. DE-AC02-05CH11231.

**Supplementary data,** including spreadsheets listing all LHE sequences and corresponding expression and cleavage values as described in the methods and results sections, are available at NAR online.

## Data Availability

The crystallographic coordinates and diffraction data corresponding to the structure of aHE31 and aHE15 have been deposited in the protein database (www.rcsb.org) and are available for immediate downloading and examination, via PDB ID codes 12FK and 36OF. Raw data (sequencing output, gels, and flow cytometric data) have been deposited and made available for public accession at the Harvard Datavase Depository (https://doi.org/10.7910/DVN/Z4WCBW).

---

|  | **aHE31** | **aHE15** |
| --- | --- | --- |
| PDB ID | 12FK | 36OF |
| **Data Collection** |  |  |
| Space group | P 1 21 1 | P 21 21 21 |
| Unit cell |  |  |
| a, b, c (Å) | 58.45, 47.37.74.74 | 53.89, 54.04, 164.83 |
| alpha, beta, gamma (°) | 90.00, 98.65, 90.00 | 90.00, 90.00, 90.00 |
| Wavelength (Å) | 1.00 | 0.98 |
| Resolution range (Å) | 47.37-2.17 (2.24-2.17) | 45.19-2.37 (2.46-2.37) |
| Unique reflections | 78378 (2215) | 36820 (732) |
| R-merge | 0.258 (1.031) | 0.089 (0.674) |
| R-meas | 0.280 (1.124) | 0.093 (0.717) |
| R-pim | 0.109 (0.440) | 0.026 (0.232) |
| CC1/2 | 0.985 (0.689) | 0.999 (0.881) |
| I/sigma(I) | 6.2 (1.9) | 19.1 (2.4) |
| Chi^2 | 1.05 (0.98) | 0.93 (0.59) |
| Multiplicity | 6.6 (6.3) | 12.1 (8.4) |
| Completeness (%) | 98.9 (96.9) | 92.7 (61.1) |
| Wilson B-factor | 21.72 | 41.17 |
| **Refinement** |  |  |
| R-work | 0.2303 | 0.2306 |
| R-free | 0.2855 | 0.2785 |
| Number of non-hydrogen atoms | 3694 | 3401 |
| macromolecules | 3391 | 3336 |
| ligands | 2 | 2 |
| water | 301 | 63 |
| Protein residues | 298 | 295 |
| RMS Bonds (Å) | 0.004 | 0.004 |
| RMS Angle (°) | 0.69 | 0.6 |
| Ramachandran favored (%) | 95.61 | 97.61 |
| Ramachandran allowed (%) | 4.39 | 2.39 |
| Ramachandran outliers (%) | 0.00 | 0.00 |
| Clashscore | 5.8 | 5.2 |
| Average B-factor | 25.13 | 49.67 |
| macromolecules | 24.12 | 45.55 |
| ligands | 12.25 | 53.83 |
| solvent | 28.89 | 41.59 |

## Supplementary Material

**Supplementary Figure S1.**
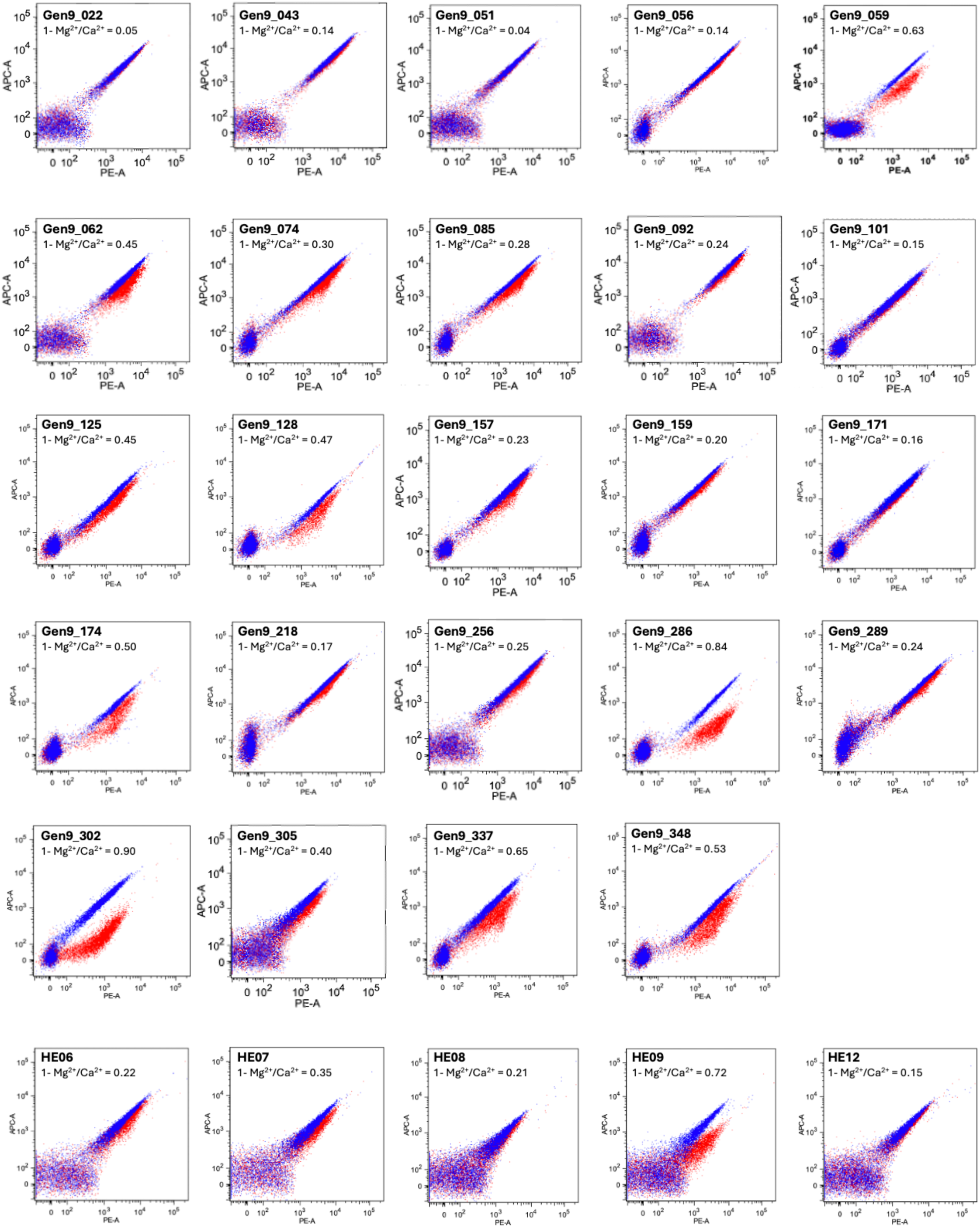
Tethered Flow Cytometric Cleavage Assay Data for 29 Active LHEs. Flow cytometric plots are shown for the 29 LHEs determined to be active against their predicted DNA target sites. Each plot is a superposition of yeast with tethered DNA target substrate in the presence of buffer with Ca^2+^ (blue, no cleavage of the tethered DNA) or in the presence of buffer with Mg^2+^ (red, cleavage of the tethered DNA). When target cleavage occurs, the cut portion of the substrate containing the A647 tag is washed away, resulting in a drop in A647 signal. If the LHE fails to release the DNA after cleavage has occurred (referred to as “end-holding”), there will be no drop in A647 signal. The quantified cleavage values (1 – Mg^2+^/Ca^2+^ Ratio) are listed for each enzyme, with a threshold activity value of greater than 0.1 used to designate active enzymes. Two of the 29 active LHEs (Gen9-022 and Gen9-051) had quantified activity values less than the 0.1 cutoff, but showed activity in the complementary non-tethered in vitro cleavage assay.

**Supplementary Figure S2.**
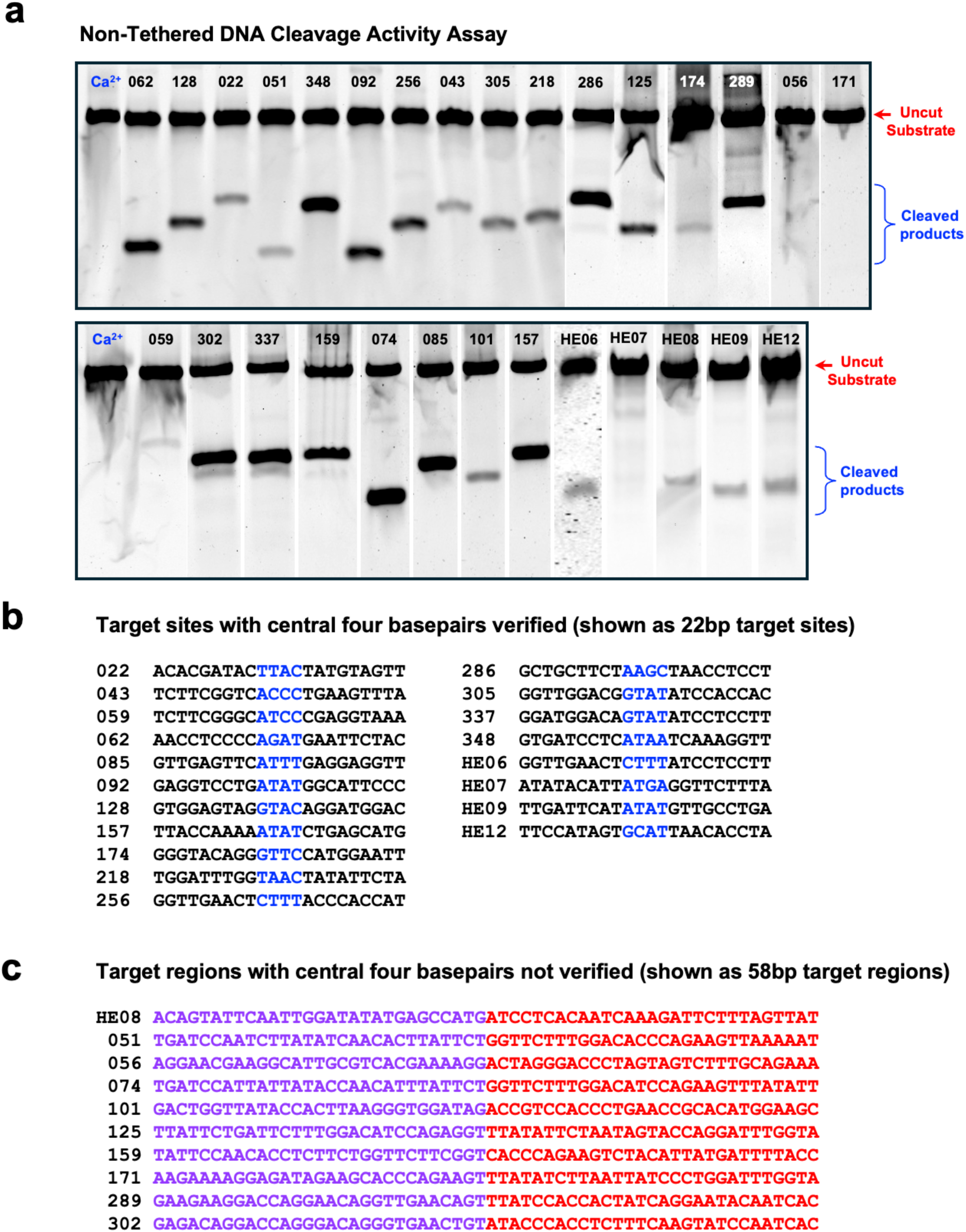
Cleavage activity in the non-tethered in vitro DNA cleavage assay for 29 active LHEs. **(A)** Gel electrophoresis of cleavage products from the non-tethered in vitro cleavage assay, imaged on a Typhoon fluorescent imager. DNA bands are visible due to the presence of the fluorescent A647 tag. Images of gels from separate experiments on multiple days are combined into one image. The red arrow indicates the location of the uncut DNA target substrates, which aligns with the lane containing the Ca^2+^ uncut control. Blue brackets indicate the location of successfully cleaved DNA fragments. The position of the cut within each 58bp target site substrate was not necessarily in the center of the DNA substrate, so the cleaved products are not all the same size. Three of the active enzymes were presumed to be weak binders (Gen9-056, Gen9-171, HE07), as they did not produce cleavage products in this assay even though they showed cleavage in the tethered flow cytometric cleavage assay. **(B)** Centered 22 base pair target sites of the active enzymes for which the precise location of cleavage on each DNA strand could be determined using run-off sequencing. The central four basepairs (which are flanked by the sites of phosphoryl hydrolysis on each individual DNA strand, producing corresponding 4-base, 3’ overhangs from those basepairs) are indicated in blue. **(C)** For those LHEs where the precise center of the DNA target sequences could not be determined (inability to isolate sufficient amounts of gel-purified products or failure to obtain clean run-off sequencing data), the sequence of the full 58 bp DNA target substrate is shown. The purple and red bases correspond to the flanking 5’ and 3’ exon sequences that were joined together to produce the predicted target site region for each candidate LHE. The site of cleavage occurs somewhere within these 58bp DNA sequences.

**Supplementary Figure S3.**
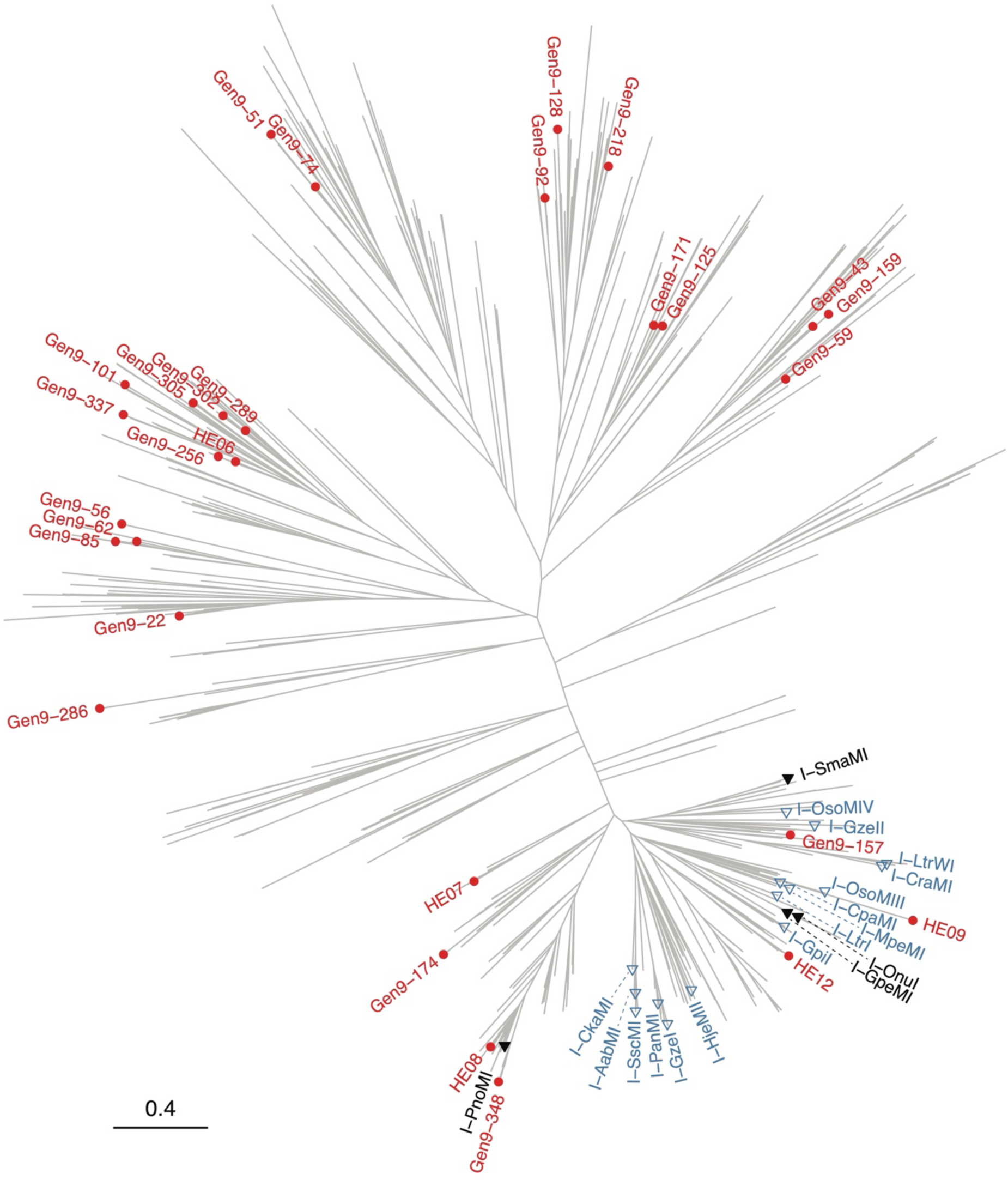
Phylogenetic tree of expanded set of LHEs. We aligned our larger 2023 dataset of 622 LHE sequences together with 11 sequences from the original experimentally tested dataset that were inadvertently missing from the 2023 set, making a total of 633 sequences. We aligned all 633 sequences using MAFFT, and removed alignment positions where >50% of sequences contained a gap. In the resulting alignment, two sequences were quite truncated and were removed before we estimated a phylogeny using PHYML, displayed using R and the ggtree package. See Methods for full details. Red labeled dots indicate the 29 newly identified active LHEs, and blue/black labeled triangles indicate 19 previously characterized LHEs, with selected sequences used in ancestral reconstruction highlighted in black.

**Supplementary Figure S4.**
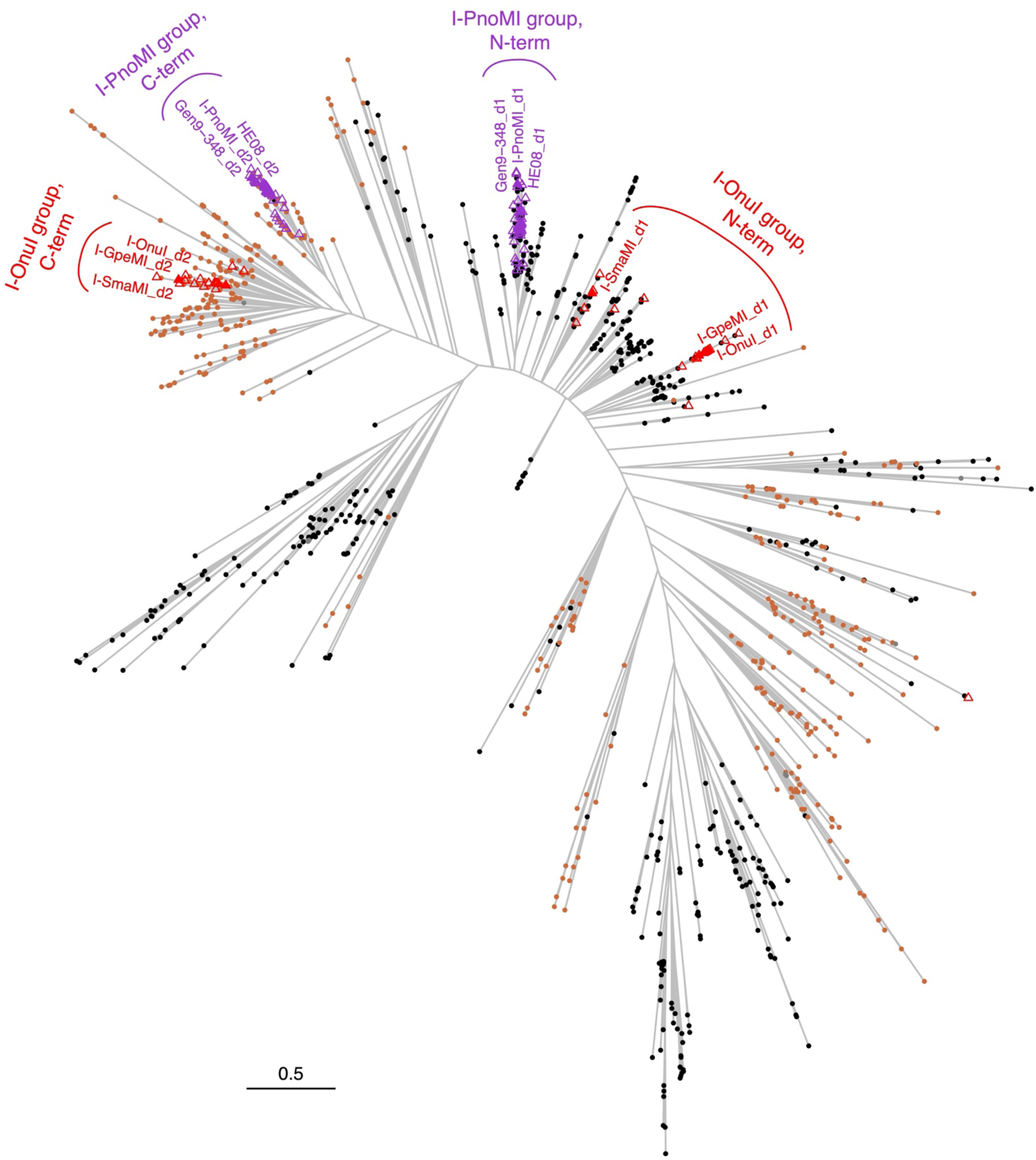
Phylogenetic tree of all individual LADLIDADG domains extracted from the dataset of 622 candidate LHEs. We used the hmmsearch tool to identify and align all matches to PFAM’s for the “LAGLIDADG_1” domain Hidden Markov Model (PF00961.22, 102aa long). Filtering to remove apparently truncated domain sequences (matching <75 aa of the domain HMM) resulted in an alignment of 1140 domains from 610 candidate LHEs. We estimated a maximum likelihood tree using PHYML and used the ggtree R package to display the tree with annotations. Almost all sequences contained two LAGLIDADG domain matches: black dots at tips of the tree are N-terminal domains, and brown dots are C-terminal domains. The scale bar indicates number of amino acid substitutions per site. We used this tree to select candidate LHEs for ancestral reconstructions for each of two sequence groups of interest: (a) I-PnoMI, Gen9-348 and HE08 (purple) and (b) I-OnuI and I-GpeMI (red). For each of the two sequence domains in each of the two groups, we located the focal members (purple/red filled triangles and text labels, with “_d1” and “_d2” indicating the N- and C-terminal domains, respectively) and identified their most recent common ancestral node. We stepped back three nodes deeper in the tree to recruit some outgroup sequences, and extracted a list of all descendants of that deeper node. We then merged the lists from the two domains in each group, and used the resulting merged sequence lists (purple/red triangles) as the bases for ancestral reconstructions. Complete details are provided in the Methods section. The selected group members for the I-PnoMI group (purple) are found very near one another on the tree for both the N-and C-terminal domains, as expected. However, for the I-OnuI group, only the C-terminal domains group closely, whereas the N-terminal domains are more scattered in the tree. This pattern is likely due to incongruent evolutionary histories between the two domains, perhaps due to recombination/gene conversion: exploring that question further is beyond the scope of this study.

**Supplementary Figure S5.**
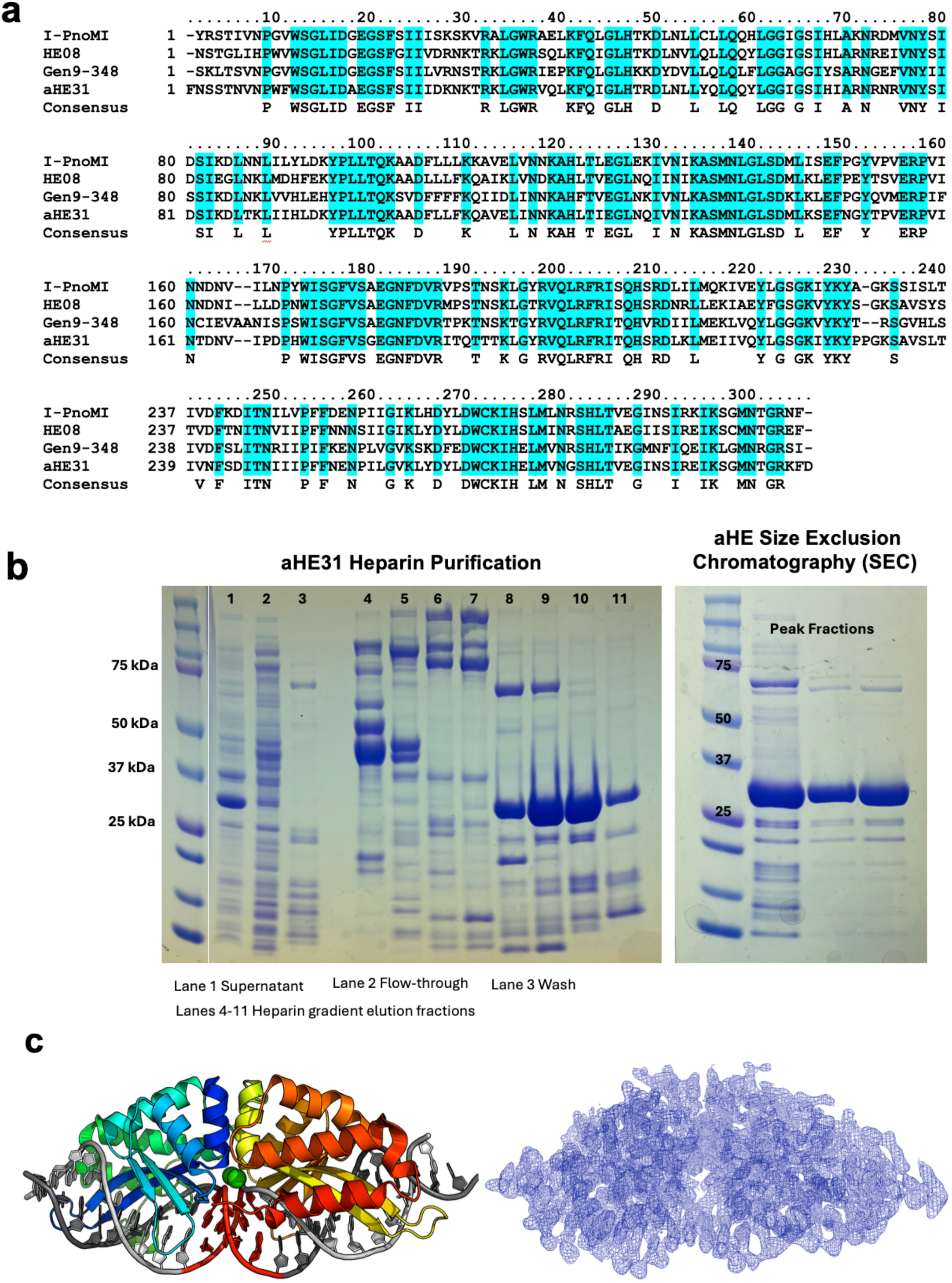
Amino acid sequence alignment of the characterized I-PnoMI clade enzymes with reconstructed ancestor aHE31 and purification of recombinant aHE31 protein. **(A)** Fully conserved positions are highlighted in cyan, with the conserved sidechains listed below the alignment. **(B)** Recombinant aHE31 protein was purified using a two-step protocol, including heparin purification followed by size exclusion chromatography (SEC). The contents of lanes on the gel from the heparin purification are listed below the gel, and the aHE31 protein is expected at a molecular weight of 34.7 kDa. **(C)** The crystal structure of the aHE31 ancestral reconstruction, bound to the I-PnoMI DNA target site, was solved to 2.17 Å resolution. See **Table 1** for data and refinement statistics.

**Supplementary Figure S6.**
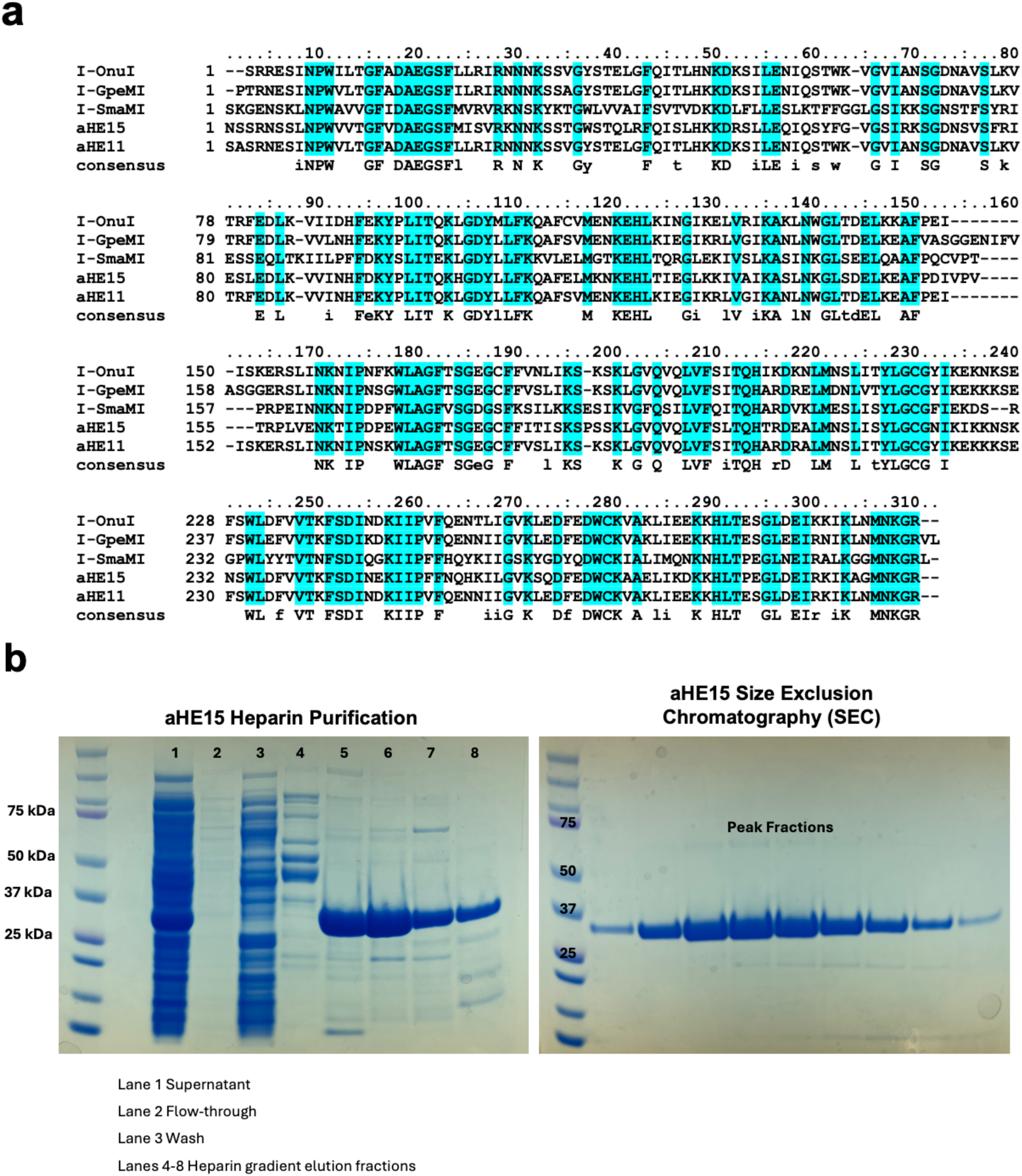
Amino acid sequence alignment of the characterized I-OnuI clade enzymes with reconstructed ancestors aHE11 and aHE15 and purification of recombinant aHE15 protein. **(A)** Fully conserved positions are highlighted in cyan, with the conserved sidechains listed below the alignment. **(B)** Recombinant aHE15 protein was purified using a two-step protocol, including heparin purification followed by size exclusion chromatography (SEC). The contents of lanes on the gel from the heparin purification are listed below the gel, and the aHE15 protein is expected at a molecular weight of 33.5 kDa.

**Supplementary Figure S7.**
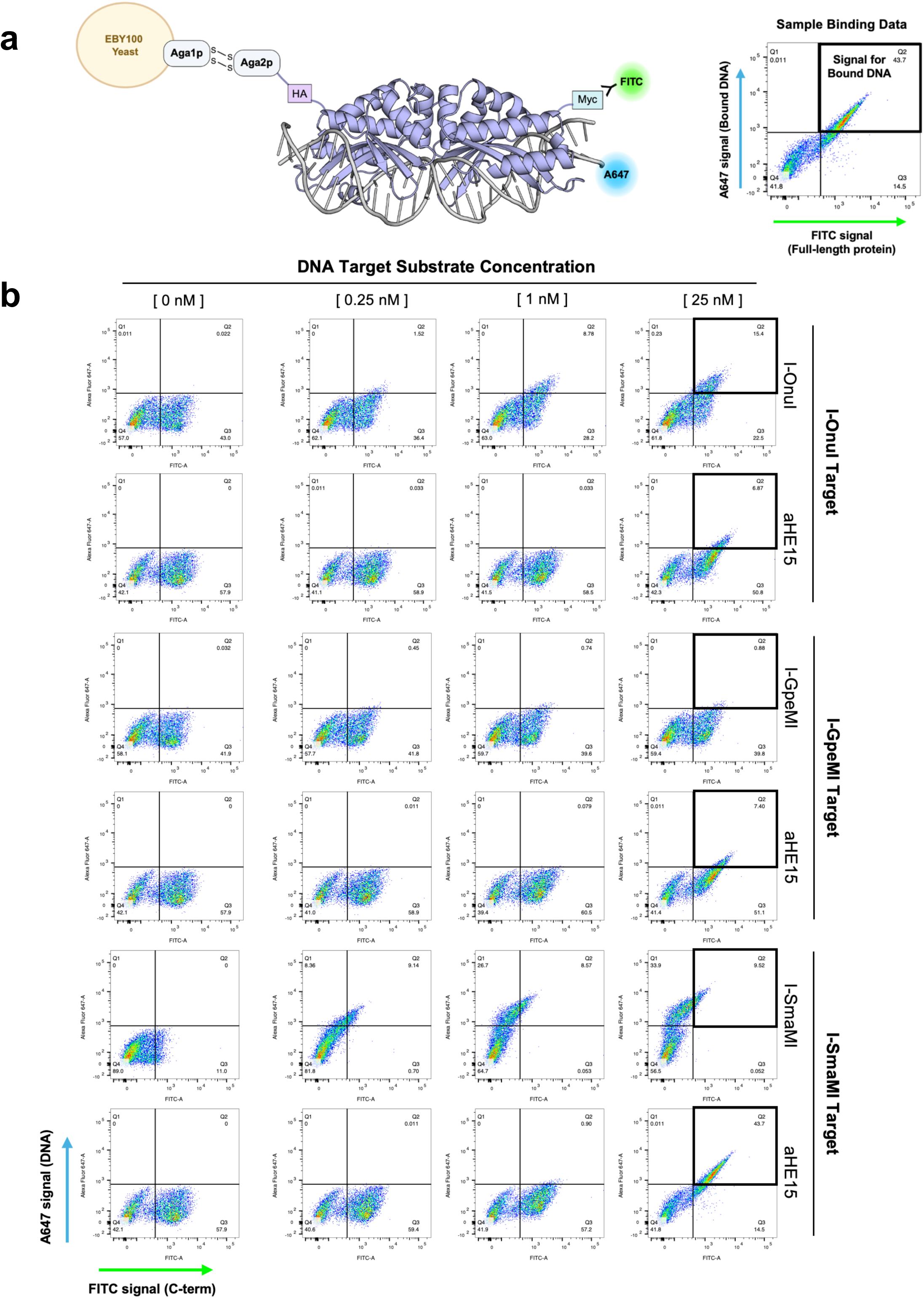
Flow cytometric binding assay to assess the ability of ancestor aHE15 to bind the DNA target sequences of related modern-day enzymes. **(A)** Schematic of the flow cytometric DNA binding assay showing an example of strong binding. Surface-expressed LHE is stained with anti-Myc-FITC to track full-length protein, and a double-stranded A647-labeled DNA target substrate is provided free in solution (no tethering). Binding of the target substrate is indicated by the presence of A647 signal in the upper portion of flow plot with FITC signal (full length protein) plotted on the x-axis and A647 signal (on the substrate DNA) on the y-axis. A bold black box highlights the portion of the flow plot where evidence of binding can be observed. **(B)** Surface-expressed aHE15 was analyzed for binding of A647-labeled DNA substrates containing the target site sequences of I-Onu, I-GpeMI, and I-SmaMI and compared to the ability of each modern-day enzyme to bind its own target sequence. All surface-expressed LHEs were stained with anti-Myc-FITC to track the presence of full-length enzyme (FITC signal, x-axis), and incubated with A647-labeled DNA target substrate ranging in concentration from 0 nM up to 25 nM (A647 signal, y-axis).

**Supplementary Figure S8:**
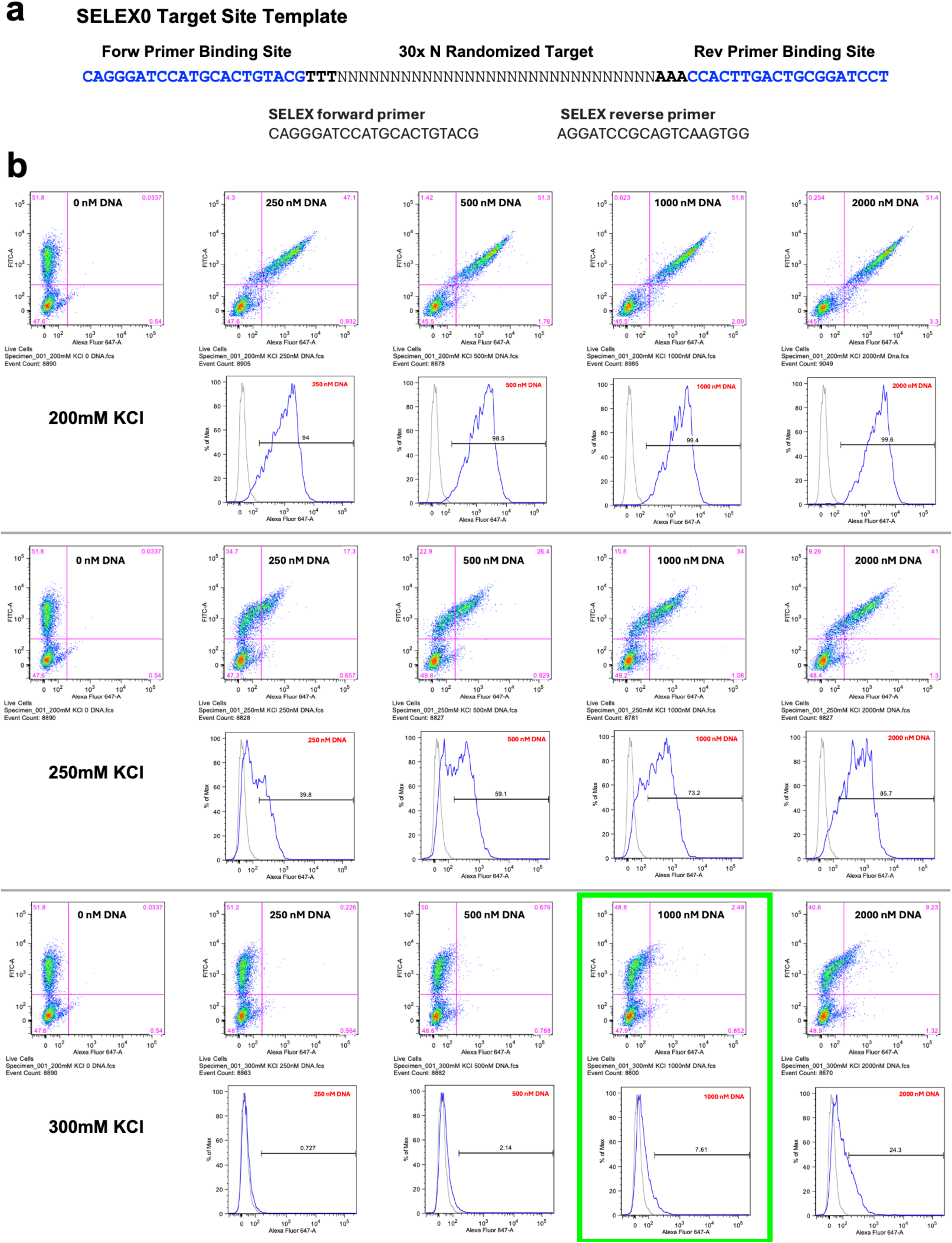
DNA target substrate design and parameter optimization for the binding-based SELEX protocol. **(A)** The SELEX0 DNA target substrate library was generated using a single stranded template consisting of 30 basepairs of fully randomized DNA sequence (“30xN”) flanked by 24 bp forward and 22 bp reverse primer sequences. Three thymidine bases preceding and three adenine bases following the 30xN sequence are designed to discourage high-affinity binding to the constant regions of the substrate. Double-stranded DNA target substrate was generated by PCR amplification with forward and reverse SELEX primers. Fluorescently labeled DNA target substrate was generated using an A647-labeled reverse primer. **(B)** The concentration of KCl in the binding buffer and total DNA target substrate in the reaction both needed to be optimized to allow for approximately 5% overall binding of the SELEX0 target library. aHE15 was expressed on the surface of yeast and assayed for binding using an A647-labeled SELEX0 substrate. KCl in the binding buffer was assayed at 200 mM, 250 mM, and 300 mM concentrations in combination with total SELEX0 substrate concentrations ranging from 250 nM – 2000 nM. The final parameters selected for the SELEX experiment included 300 mM KCl and 1000 nM SELEX0 target site library substrate, which allowed for binding of approximately 7.6% of the SELEX0 library (bold green box).

**Supplementary Figure S9.**
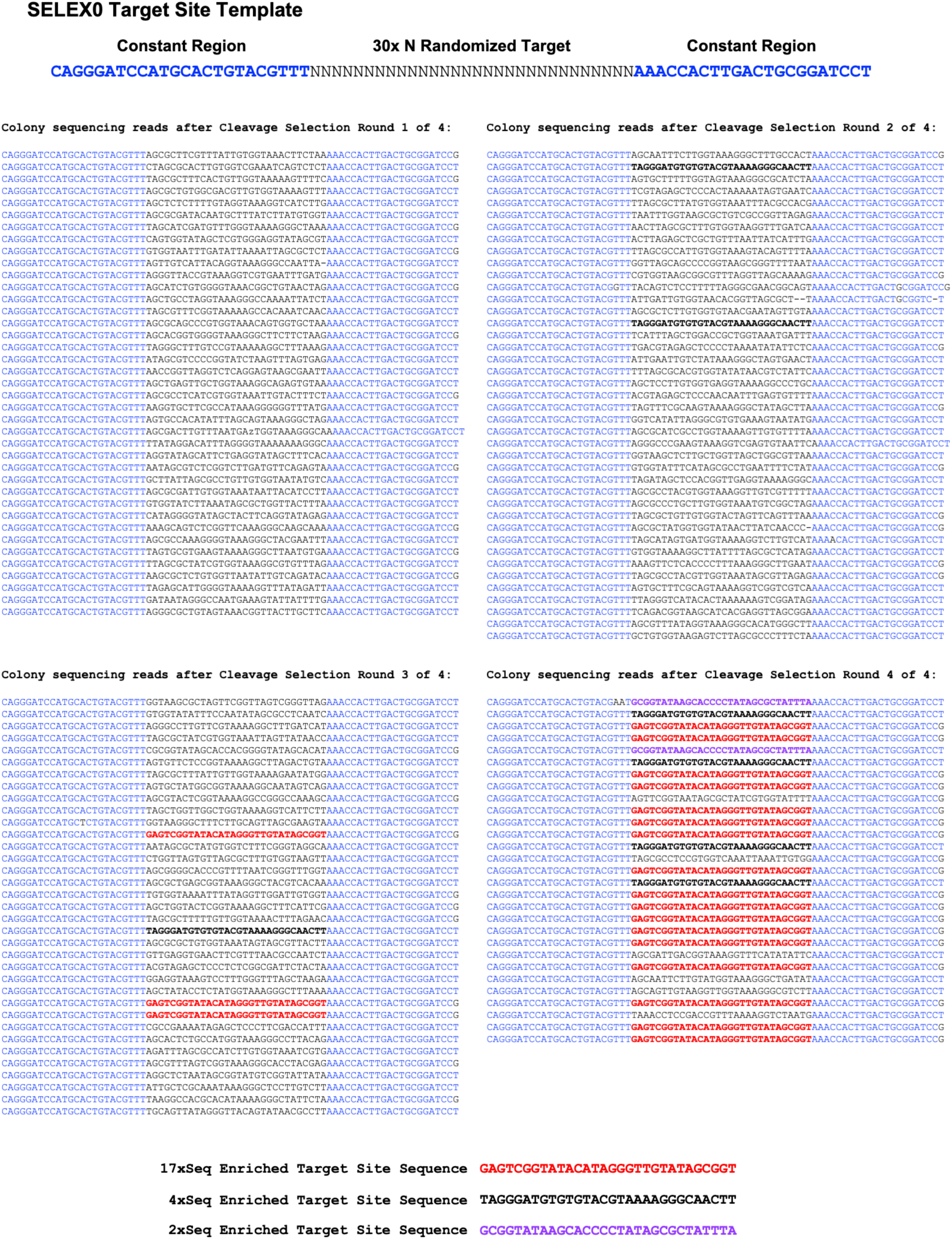
Sequencing results from 4 rounds of cleavage selection. 48 colonies were sequenced after each round of the cleavage selection to look for enrichment of sequences cleavable by the aHE15 enzyme. Failed or ambiguous sequence reads are not listed here. The constant regions of the DNA target substrate mark the boundaries of the 30 bp target sequence under analysis and are colored with blue text. After four rounds of cleavage (bottom right), there were three different target sequences that were enriched to some degree, with the “2xSeq” target present two times (purple bold text), the “4xSeq” target present four times (black bold text), and the “17xSeq” target present in 17 of the 29 good sequence reads (red bold text). The 17xSeq target sequence was also present 3 times in the colonies sequenced after Round 3, and the 4xSeq target was present once after Round 3 and two times after Round 2.

**Supplementary Figure S10.**
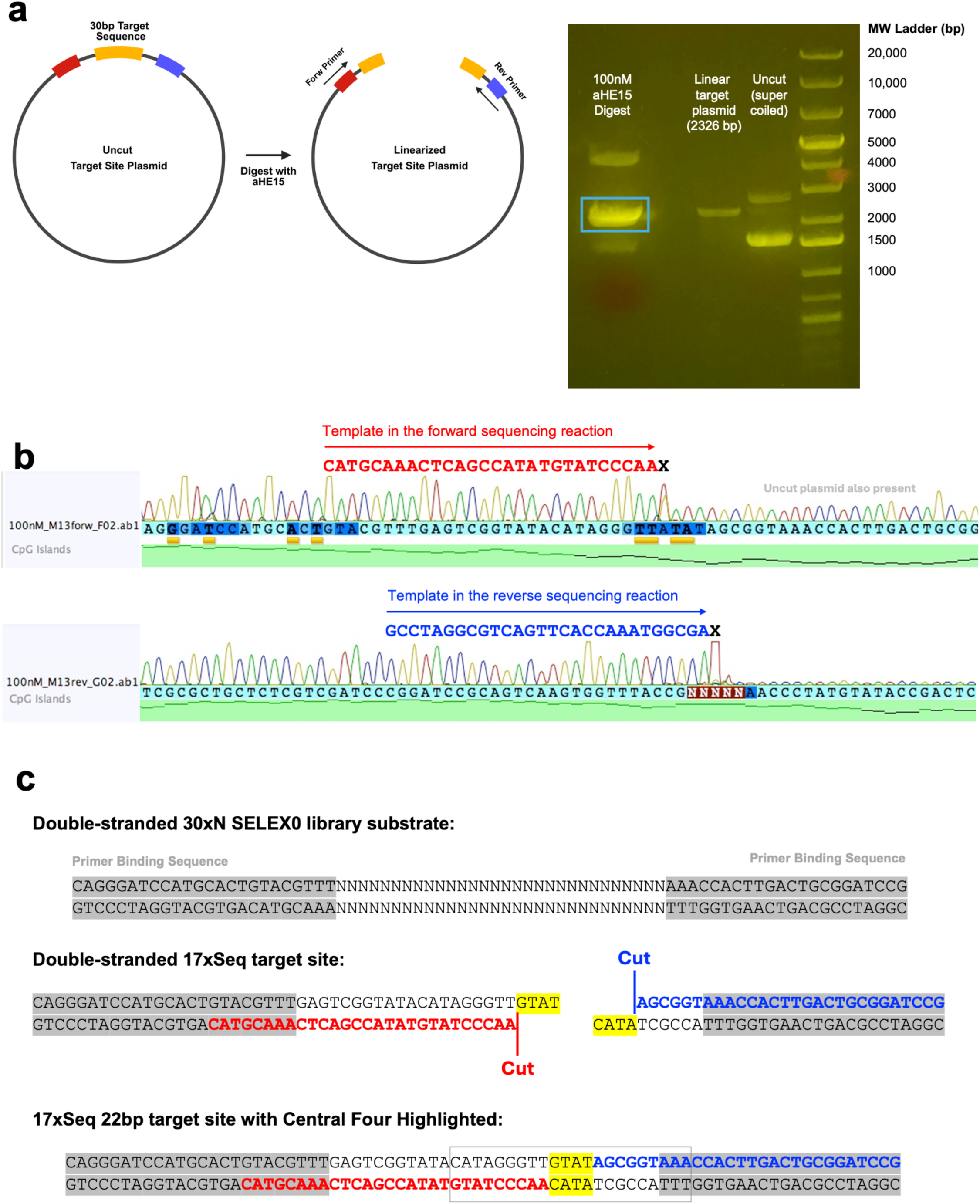
Run-off sequencing to determine the center of the 17xSeq cleavable target sequence for aHE15. **(A)** Schematic representation of the run-off sequencing experiment. The 17xSeq target site plasmid contains the 30 bp target sequence at a position that could be sequenced by both forward and reverse sequencing primers. Recombinant purified aHE15 protein was used to digest the intact 17xSeq target site plasmid, and the linearized product (blue box in the gel) was extracted from an agarose gel and purified. A PvuI-linearized plasmid control was run alongside uncut/supercoiled plasmid in the gel to help determine the correct location of the linearized product for extraction. **(B)** Both forward and reverse primers were used to sequence the purified linear product in two separate Sanger sequencing reactions. The Taq polymerase in the BigDye sequencing master mix (Thermo Fisher Scientific) produced a large adenine peak in the sequencing chromatogram when it reached the end of the cleaved DNA template strand. The location of this large A peak was used to identify the precise location where aHE15 cleaved each strand of the DNA target. The sequencing chromatogram readout represents the complement of the actual template in the sequencing reaction, so the complementary sequence of the target site plasmid template in the reaction is listed above the chromatogram for each read. The sequence of the template in the forward sequencing reaction is listed in red text and the template in the reverse sequencing reaction is listed in blue text. A black **X** indicates the position in the DNA sequence of the cut made by the aHE15 enzyme, as indicated by the large “A” peak in the chromatogram. **(C)** The double-stranded sequence of the 30xN SELEX0 library target site substrate is listed for reference, followed by the double-stranded 17xSeq target sequence identified after round 4 of the cleavage selection. The locations of the observed cuts on each strand of the target sequence (identified by the two run-off sequencing reactions shown above) are marked with vertical red and blue lines, and spaces are introduced at the position of each cut to show the 4-base, 3’ overhangs generated by the aHE15 enzyme. Finally, the 22bp centered 17xSeq target is outlined with a rectangular box.

**Supplementary Figure S11.**
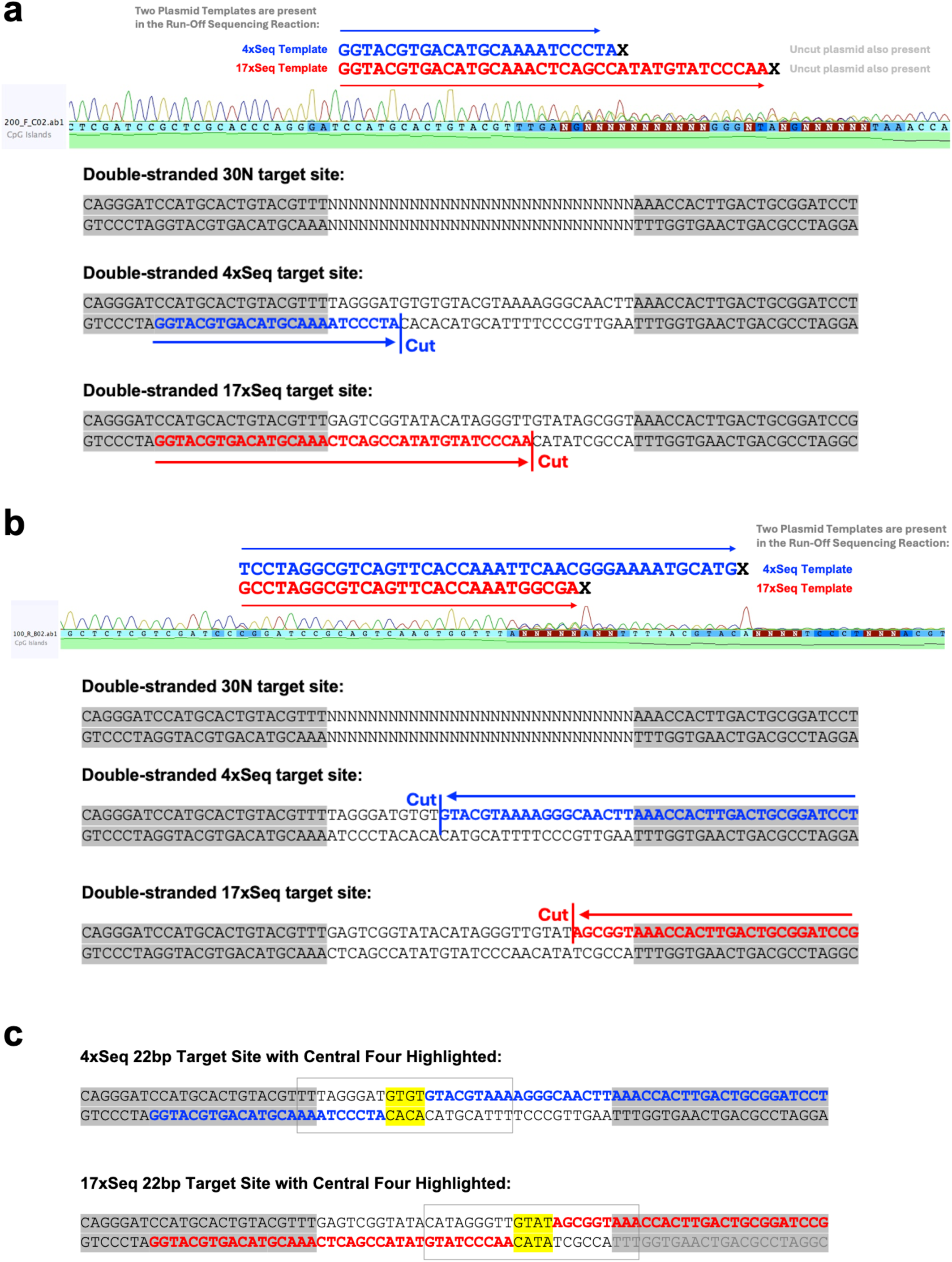
Run-off sequencing to determine the center of the 4xSeq cleavable target sequence for aHE15. **(A)** A forward run-off sequencing reaction with linearized 4xSeq target site plasmid as the template revealed that the prep actually contained two different plasmids containing the target site plasmids for both the 4xSeq and 17xSeq sequences. It was possible to identify the chromatogram peaks corresponding to each of the two separate target sequences (4xSeq target in blue text and 17xSeq target in red text), as well as two separate large “A” peaks to represent two cleavage locations. There was also signal for uncut plasmid leftover in the digest reaction, as indicated by the presence of additional smaller peaks in the sequencing chromatogram after the location of cleavage. **(B)** The reverse run-off sequencing reaction also contained chromatograms for both the 4xSeq (blue text) and 17xSeq (red text) targets, as well as two separate large “A” peaks to designate the location of cleavage. **(C)** The run-off sequencing reactions allowed for the identification of the centered 22 bp target sequences for both the 4xSeq (blue) and 17xSeq (red) target sites, as outlined by rectangular boxes. The centered 22 bp 4xSeq target includes two of the flanking thymidines from the constant region of the SELEX target site template, and the centered 22bp 17x Seq target includes all three of the flanking adenines from the constant region. The center of the 17xSeq target site identified here matched and confirmed the data presented in **Supplementary Figure S10**.

**Supplementary Figure S12.**
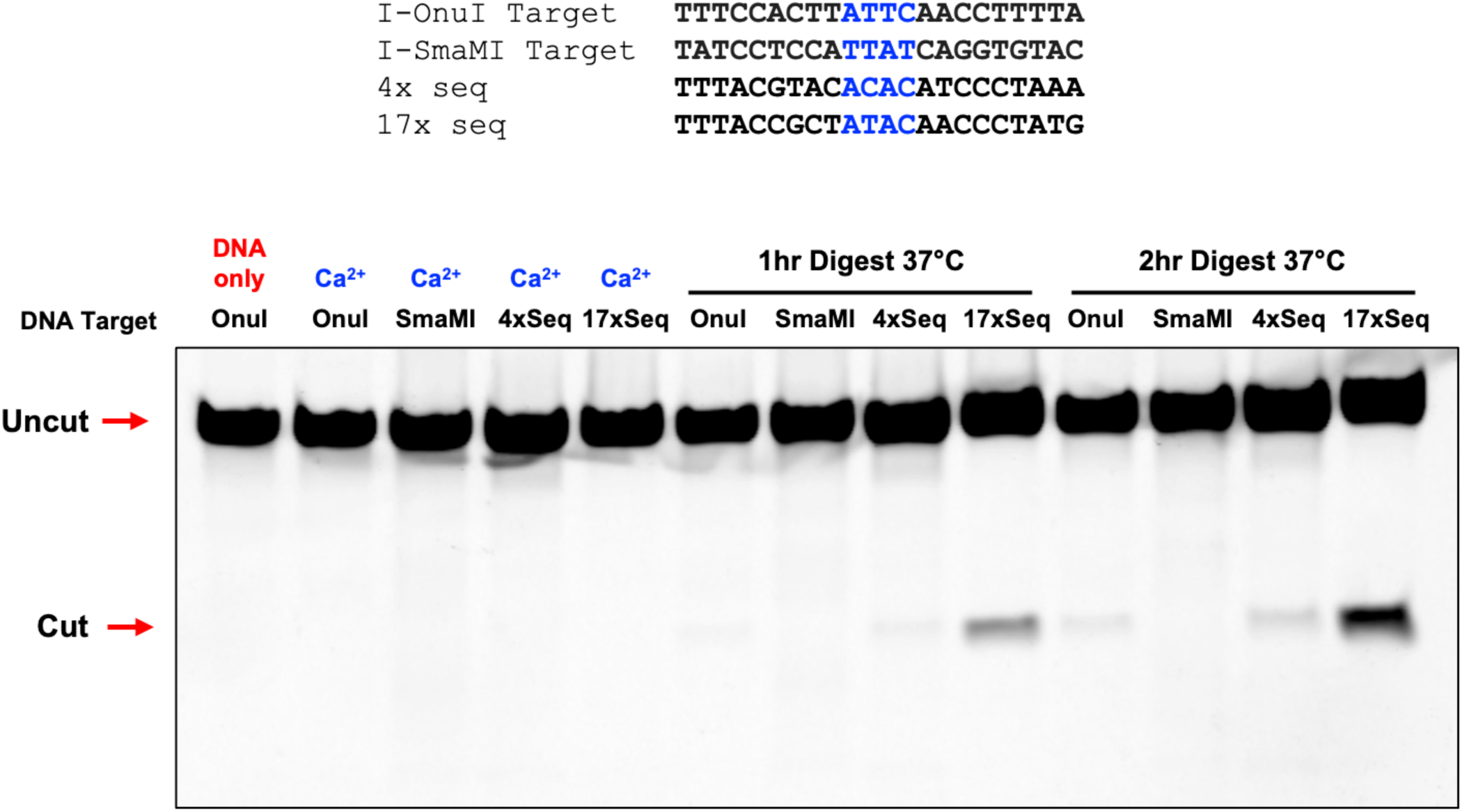
Cleavage activity of aHE15 vs. the two most highly enriched sequences from the cleavage selection screen. Purified recombinant aHE15 was used to digest A647-labeled DNA substrates containing four different target sequences: I-OnuI, I-SmaMI, and both the 4xSeq and 17xSeq targets that were enriched after four rounds of the cleavage selection screen. The 22bp sequences for the 17xSeq and 4xSeq targets are shown as the reverse complement of the sequences presented in Figures S9 - S11, as this orientation lines up best with the verified I-OnuI target sequence. The sequences of the 17xSeq and 4xSeq targets are identical at 14 out of 22 positions. Cleavage products were separated by electrophoresis on an acrylamide gel and visualized by the presence of the A647 tag using a Typhoon fluorescence imager. Controls included a DNA-only lane (no enzyme) and digests performed in the presence of CaCl_2_ (Ca^2+^ lanes) to indicate the position of uncut DNA substrate. The locations of cut and uncut DNA substrates are indicated on the gel with red arrows.

**Supplementary Figure S13.**
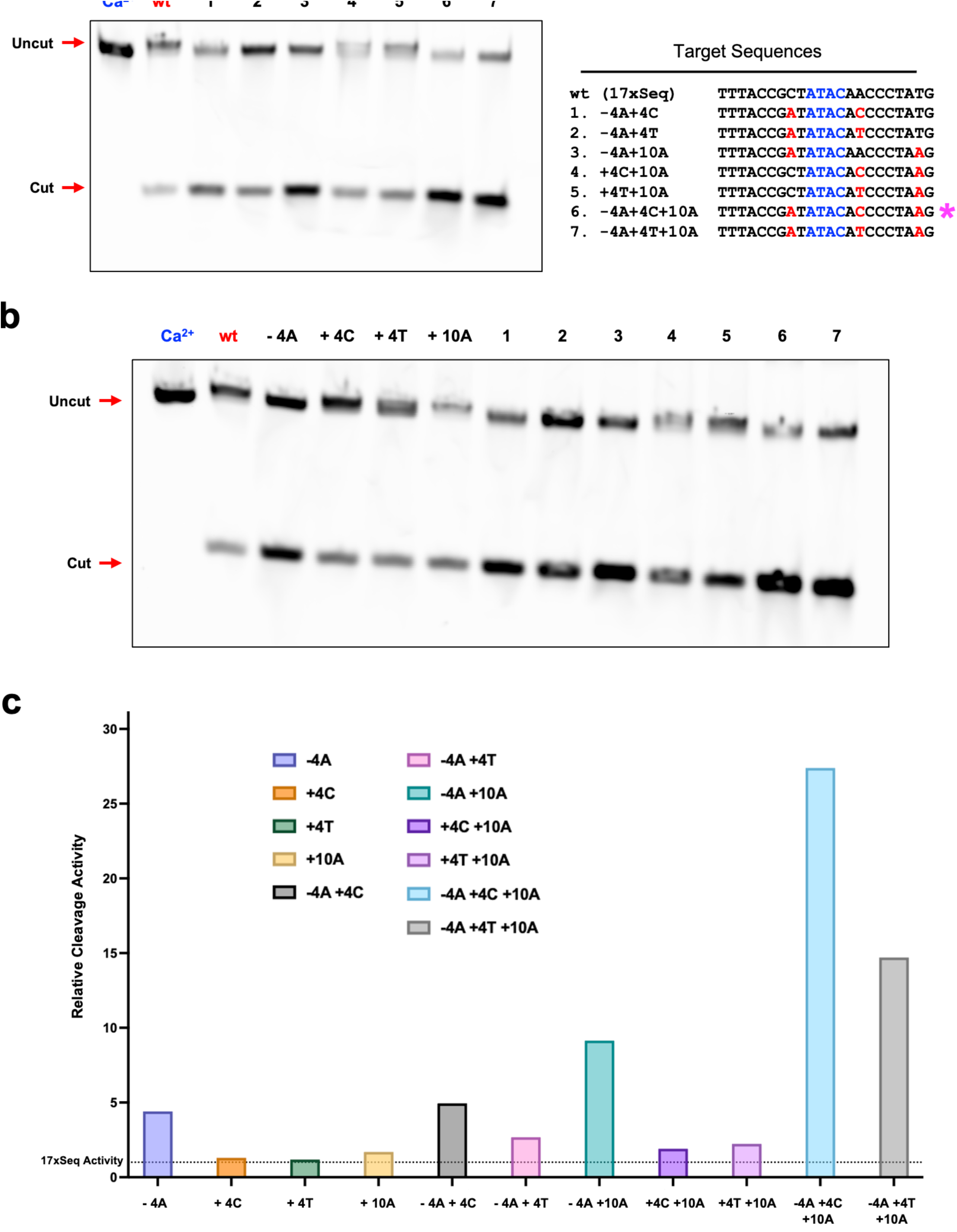
Gel of in vitro digests used to identify an optimized alternative target sequence for aHE15. **(A)** Image of the acrylamide gel with cleavage products from *in vitro* digests of the seven combinatorial alternative target site sequences. The far-left lane contains an uncut control from a reaction run with CaCl_2_ (blue Ca^2+^ label), and the location of cut and uncut products are marked with red arrows. The full 22bp DNA sequences of the seven alternative target sites are listed to the right of the gel, with each single basepair change away from the original 17xSeq sequence highlighted in red. A magenta asterisk designates the target cleaved best by the aHE15 enzyme. **(B)** Acrylamide gel from a repeat of the experiment comparing the cleavage of all four single off-targets side-by-side with the original 17xSeqtarget and the seven combinatorial targets. A Ca^2+^ lane serves as an uncut control, and the location of cut and uncut products are marked with red arrows. **(C)** Quantification of the cleavage products from the gel shown in **Supplementary Figure S13b** with bars representing relative cleavage activity compared to the cleavage of the original target sequence (17xSeq). A dotted line designates the level of cleavage of the original 17xSeq target (a relative cleavage activity of 1.0.)

