## Supplementary sequence information for "The activity, divergence, and evolutionary degradation of modern-day homing endonucleases and their reconstructed ancestors": suppFiles_trees_README.md

Files related to Figure 3: 

`fig3a_alignmentForTree_n390.fa` is a fasta file of the degapped and filtered sequence alignment used as input for the PHYML tree shown in Figure 3A. 

`fig3a_sequenceInfoTable_n390.csv` provides more information about each sequence, including the full-length unaligned version of each sequence, is provided in the file 


`Fig3B_PnoMI_ancReconstr_alignment_2026_07_10.fa` and `Fig3C_OnuI_ancReconstr_alignment_2026_07_10.fa`. Fasta format sequence files containing BAli-Phy's summary alignments of extant sequences with selected ancestral nodes, as shown in Figures 3B and 3C. For the I-PnoMI clade, we used the sequence of aHE31 exactly as output by BAli-Phy.  For the I-OnuI clade, we noticed that in several regions of the alignment, BAli-Phy predicted amino acid residues in ancestral sequences where gap characters are much more parsimonious, and we edited the predicted ancestral sequences accordingly. The I-OnuI clade alignment file therefores contains edited and unedited versions of the sequences at ancestral nodes aHE11 and aHE15.


Files related to Supp Figure 3: 

`suppFigS3_alignmentForTree_n631.fa` is a fasta file of the degapped and filtered sequence alignment used as input for the PHYML tree shown in Supp Figure 3. Sequences identifiers in this file are as provided by author BLS, and alternative identifiers that match those used elsewhere in this study can be found in the associated csv information table (below). 

`suppFigS3_sequenceInfoTable_n633.csv` provides more information about each sequence, including the full-length unaligned version of each sequence. It also includes information about the two sequences that were removed from the alignment because they were too short after degapping.
 


Files related to Supp Figure 4: 

`suppFigS4_domainAlignmentForTree_n1140.fa` is a fasta file of the degapped and filtered sequence alignment used as input for the PHYML tree shown in Supp Figure 4. Sequence identifiers in this file are of the form "A/B-C", where A is the full-length sequence identifier (found in file `suppFigS3_sequenceInfoTable_n633.csv`), and B-C are the amino acid coordinates of the Hidden Markov Model's LADGLIDADG domain match - the region used for these alignments. Note that residues that do not align to the HMM are not included in this alignment.
